# Social modulation of oogenesis and egg-laying in *Drosophila melanogaster*

**DOI:** 10.1101/2021.09.13.460109

**Authors:** Tiphaine P. M. Bailly, Philip Kohlmeier, Rampal S. Etienne, Bregje Wertheim, Jean-Christophe Billeter

**Affiliations:** Groningen Institute for Evolutionary Life Sciences, University of Groningen, Groningen, The Netherlands

**Keywords:** social environment, *Drosophila melanogaster*, sociality, light, vision, oogenesis, egg-laying

## Abstract

Being part of a group facilitates cooperation between group members, but also creates competition for limited resources. This conundrum is problematic for gravid females who benefit from being in a group, but whose future offspring may struggle for access to nutrition in larger groups. Females should thus modulate their reproductive output depending on their social context. Although social-context dependent modulation of reproduction is documented in a broad range of species, its underlying mechanisms and functions are poorly understood. In the fruit fly *Drosophila melanogaster,* females actively attract conspecifics to lay eggs on the same resources, generating groups in which individuals may cooperate or compete. The tractability of the genetics of this species allows dissecting the mechanisms underlying physiological adaptation to their social context.

Here, we show that females produce eggs increasingly faster as group size increases. By laying eggs faster in group than alone, females appear to reduce competition between offspring and increase their likelihood of survival. In addition, females in a group lay their eggs during the light phase of the day, while isolated females lay them during the night. We show that responses to the presence of others are determined by vision through the motion detection pathway and that flies from any sex, mating status or species can trigger these responses. The mechanisms of this modulation of egg-laying by group is connected to a lifting of the inhibition of light on oogenesis and egg-laying by stimulating hormonal pathways involving juvenile hormone. Because modulation of reproduction by social context is a hallmark of animals with higher levels of sociality, our findings represent a protosocial mechanism in a species considered solitary that may have been the target of selection for the evolution of more complex social systems.

## Introduction

Being in a group can benefit the individual by reducing predation risk (Chapman and Reiss, 1999), improved territory defense (Sakata and Katayama, 2001), enhanced protection from environmental conditions by increasing ambient humidity and reducing desiccation risk (Howe, 1962a), facilitated reproduction and increased offspring survival (Allee, 1927a; Wertheim et al., 2002b). However, individuals in groups must also compete for access to mates, feeding territories, and for raising offspring on shared resources (Clutton-Brock and Huchard, 2013; Clutton-Brock, 1991; Stockley and Bro-Jørgensen, 2011). Reproductive competition among females is observed in a range of species, leading to delayed sexual maturation, spontaneous abortions and early offspring mortality (DeLong, 1978; Dunbar R. I. M and Dunbar E. P., 1977; Laws, 1929; Peyser MR et al., 1973; Rowell, 1970; Wasser and Barash, 1983). This competition obviously limits the scope for cooperation. For this reason, mechanisms suppressing inter-individual competition are thought to be a major tenet of the evolution of sociality (Frank, 2003). Mechanisms that suppress reproductive competition are indeed key to eusociality, where division of labour allows only one or few female(s) to be reproductively active while the other are functionally infertile and help the reproductive success of the reproducing individuals through foraging, guarding and nursing (Wilson, 1971).

Although the actual fertilization of the oocyte requires only a male and a female, animals ranging from mammals to insects and across a gradient of social complexity modulate their reproduction depending on social context. For instance, female rats experience ovarian synchrony with other female group members (McClintock, 1984; Schank and McClintock, 1997). In fish, females reduce egg size with increasing group member number, thus adjusting investment in offspring to group size (Taborsky et al., 2007). In eusocial insects, workers oocyte size increases with the number of workers in the colony, while it decreases as the number of larvae increases (Mohammedi et al., 1998; Oldroyd et al., 2001; Traynor et al., 2014; Ulrich et al., 2016; Villalta et al., 2015). In gregarious cockroaches, grouped females display advanced oocyte maturation and increased oviposition compared to isolated females (Crall et al., 2016; Uzsak and Schal, 2012; Uzsák and Schal, 2013). Stimulation of reproduction by the presence of others is also documented in a couple of insect species considered solitary, such as in *Rhagoletis pomonella* flies, where females lay more eggs in a group than when alone (Prokopy and Reynolds, 1998) and in blowflies, where egg production and oviposition exhibit positive density dependence (Davies, 1998). Despite widespread documentation of social modulation of reproduction, the function and mechanisms of such phenomena often remain elusive, but for a few examples. The queen pheromones in bees inhibit ovary development in workers, suppressing reproductive competition within a nest (Hoover et al., 2003; Le Conte et al., 2001). In gregarious cockroaches, social facilitation of reproduction is regulated by increased juvenile hormone biosynthesis which accelerates female oogenesis (Gadot et al., 1989; Uzsak and Schal, 2012). Hormones and pheromones appear to be key to social modulation of reproduction. To better understand the widespread -but poorly understood-phenomenon of social modulation of reproduction, a genetically and experimentally tractable organism capabale of such modeulation would allow investigating both the underlying behavioral and physiological mechanisms.

*Drosophila melanogaster* displays modulation of reproduction in groups and is an experimentally tractable and ideal organism for dissecting the mechanisms underpinning behaviour and reproduction. Indeed, several studies have shown that female reproduction is modulated by the presence of others in this species. For instance, females mate more frequently in the presence of more numerous and diverse males (Billeter et al., 2012; Gorter et al., 2016; Krupp et al., 2008) and they eject sperm and remate faster when they perceive the presence of other females that are seen as competitors (Laturney and Billeter, 2016). Mated females actively form groups by attracting other females via pheromones to an egg-laying substrate (Amrein, 2004; Bartelt et al., 1985; Billeter and Levine, 2015; Duménil et al., 2016a; Wertheim et al., 2002a). The function of communal egg-laying appears cooperative as it results in a greater amount of larvae, with better survival because they can fend off the growth of fungi (Trienens et al., 2017; Wertheim et al., 2002c). However, aggregating on the same substrate also has costs as larvae compete for food, culminating in cannibalism when resources are exhausted (Allee, 1927a; Courchamp et al., 1999; Etienne et al., 2002a; Narasimha et al., 2019; Stephens and Sutherland, 1999; Vijendravarma et al., 2013a; Wertheim et al., 2002b). These costs and benefits predict the existence of mechanisms that allow females to adjust the number of eggs they lay on the substrate to the number of females and offspring already present on it.

Here, we show that grouped females advance the onset of egg-laying while isolated females wait and solely lay eggs during the night. We show that start-time of egg-laying is density-dependent and that group detection is determined by vision through the motion detection pathway. We demonstrate that the presence of others trumps light conditions resulting in females laying eggs during the day, while isolated females normally retain their eggs until darkness. We further find that the presence of a group lifts the inhibition of light on oogenesis and egg-laying by stimulating hormonal pathway involving juvenile hormone. Our study therefore demonstrates social modulation of oogenesis and egg-laying in *Drosophila*, and reveals which fundamental behavioural and physiological aspects of reproduction are modulated by the presence of others.

## Results

### Females enhance offspring survival by laying eggs faster in a group than when alone

To test the hypothesis that individual females modulate their egg-laying behaviour in response to the presence of others, we designed an assay in which we could monitor egg-laying behaviour of individual females in different social contexts. A focal female was placed on an egg-laying site, alone or in the presence of a varying number of mated females immediately after mating, and we studied egg-laying behaviour through analysis of time-lapse images taken every 15 minutes. To discriminate eggs laid by the focal female from those laid by group members, we used group members whose eggs express green or red fluorescent protein. We found that focal females took approximately 3 hours longer to lay their first egg when alone than when grouped (Figure 1A). Timing of egg-laying was gradually reduced with increasing group size, to an average minimum of 1 hour after mating as group size increased to 50 females (Figure 1B). This advancement of egg-laying did not affect the overall reproductive output as grouped females laid a similar number of eggs with similar egg quality (similar egg volume and viability) compared to isolated females in the 24h following the start of the assays (Supp. Figure S1 A-C). Social context thus modulates the onset of egg-laying in *Drosophila melanogaster*.

**Figure 1:**
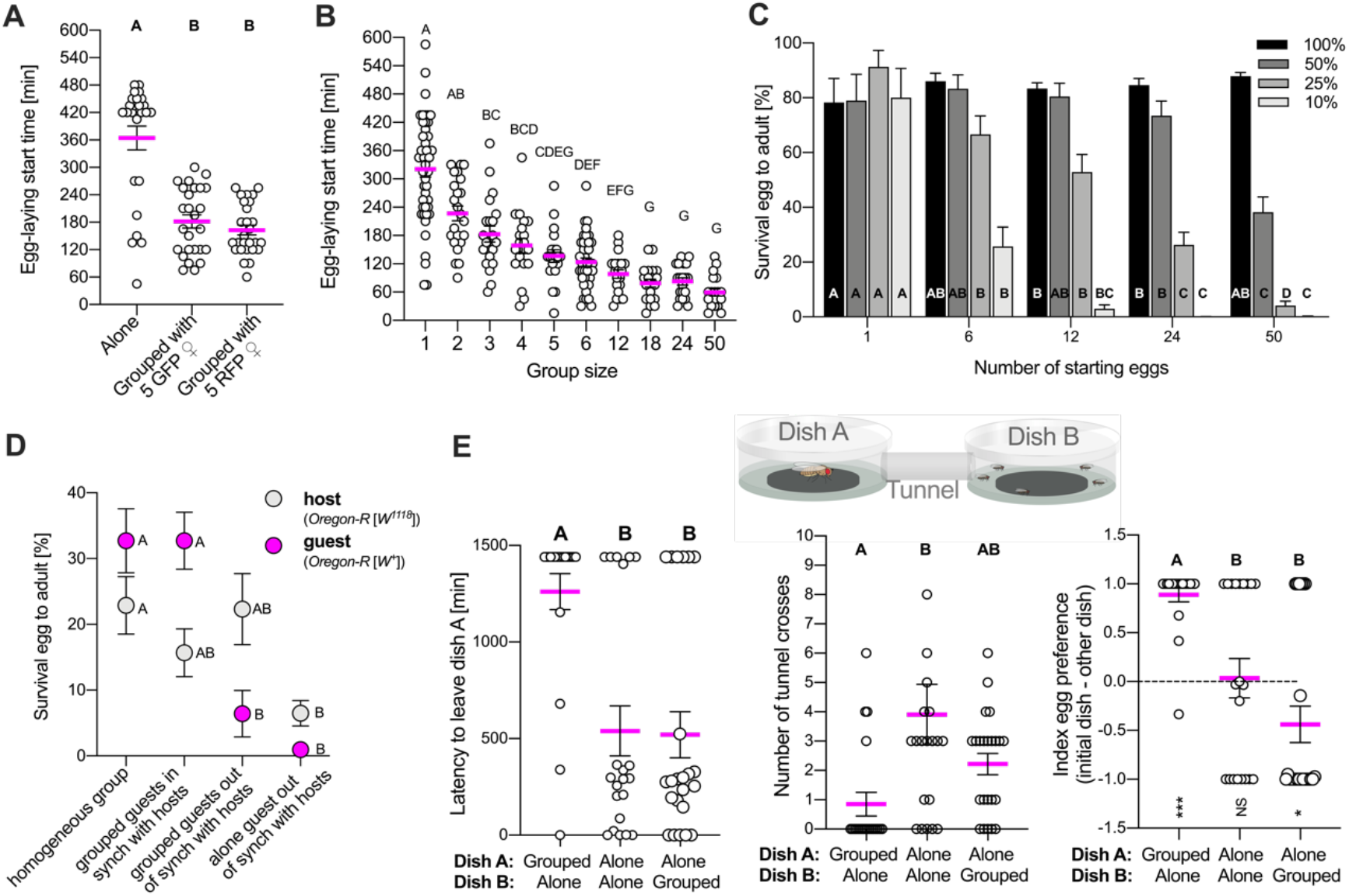
females modulate egg-laying timing depending on social context. **(A)** Onset of egg-laying of a wild-type *Oregon-R* mated female alone, grouped with 5 mated females laying green fluorescent eggs (GFP) or grouped with 5 mated females laying red fluorescent eggs (RFP). Replicates per group:25-27. **(B)** Onset of egg-laying of a wild-type *Oregon-R* mated female housed in different group sizes. Replicates per group: 20-45. **(C)** Survival of wild-type *Oregon-R* eggs housed in different group sizes in different food qualities (food containing 100%, 50%, 25% or 10% of quantities of standard food ingredients). Replicates per group: 16-23. **(D)** Survival of guest eggs from homogeneous and heterogeneous groups into viable adults when laid in synchrony or out of synchrony with host eggs. 50% food was used for the assay (food containing 50% of quantities of standard food ingredients). Replicates per group: 25-32. **(E)** Schematic representation of the choice assay. A mated female is placed in dish A alone or in a group with decapitated flies and is free to leave the dish and go to another dish to lay eggs by crossing a tunnel. Latency to leave dish A, index egg preference (positive values show preference to laying eggs in dish A than in dish B), and number of tunnel crosses were quantified on the same females Replicates per group: 19-20. Error bars indicate standard error of the mean. Different letters indicate differences between conditions. For full statistical analysis and methods, see **Supplementary Table S1.**

We hypothesised that advancing egg-laying in the presence of other females could increase offspring-survival through decreased competition for food among the offspring. From this hypothesis, two predictions can be derived: eggs are less likely to develop into a viable adult if i) competition between larvae increases and / or ii) other eggs on the same food patch are already more advanced in their development. To test these predictions, we first recorded egg survival at different egg group sizes and under different levels of food deprivation (food containing 50, 25 and 10% of quantities of standard food ingredients). In line with the prediction, we showed that the likelihood of an egg developing into a viable adult decreased with increased number of resident eggs and with the diminution of food quantities (Figure 1C). The number of eggs that females laid when they were in a group exceeded what can be sustained by a food patch, suggesting that females may lay eggs faster when in a group than alone to give a competitive advantage to their offspring who can first exploit the limited resources. Under this scenario, females need to start laying eggs on the food patch at the same time –or at least not later-than group members to increase the chances of their offspring survival. Alternatively, females in a group might modify egg composition or substrate (e.g. enzymes, faeces) leading to higher offspring performance irrespective of start-time of egg-laying. To discriminate between these two hypotheses, guest (wild-type *Oregon-R* with red eyes) and host females (*Oregon-R* with white eyes for discrimination) were used. Grouped host females laid eggs in a dish and 40 of these eggs were mixed with 10 eggs of guest females from three alternative groups: (1) grouped females who laid eggs at the same time as the hosts, (2) grouped females who laid eggs 6 hours later than hosts, (3) isolated females who laid eggs 6 hours later than hosts. The mixed groups were kept in dishes with a limited amount of food during 12 days, leading the larvae to compete for resources. We found that probabilities of egg-to-adult survival of guest eggs laid by grouped guest females synchronous to host females were higher than those of eggs laid by either grouped or isolated guests out of synchrony with the hosts (Figure 1D). This finding reveals that eggs laid at the same time as other group members ensured maximum survival. Guest egg survival in homogeneous groups was the same (about 32%) as guest egg survival synchronized with hosts in heterogeneous groups (Figure 1D). The decreased survival of guest eggs in out-of-synchronization conditions is therefore not due to a kin effect but is due to a lack of synchronization. In addition, guest eggs laid by out-of-synchronization grouped females had 6.5-fold higher chances of survival than the guest eggs laid by females that were alone (Figure 1D), suggesting social-context dependent maternal effects. Survival of hosts was about the same for each condition, except when mixed with eggs from single females (Figure 1D). This result could be explained by a larger competition in this condition leading to more cannibalism and a decrease of survival of both host and guest eggs. Taken together, these data show that females compete by laying eggs fast to give their offspring a better chance of survival.

This functional experiment indicates that females compete for egg-laying opportunities, an observation that is paradoxical because females are attracted to communal egg-laying sites (Billeter and Levine, 2015; Duménil et al., 2016a; Wertheim et al., 2002a). As females are artificially forced to be in a group in our assay, are they making the best of a bad situation or are they indeed attracted to lay eggs earlier in group than when alone when they have a choice? To answer this question, we designed a choice assay where a female can choose to lay eggs alone or in a group by crossing a tunnel (Figure 1E). Females left the starting dish more often and faster when alone than when in a group, spending more time and laying more eggs in the dish already occupied by a group of flies (Figure 1E). When having the choice between two empty dishes (alone-alone), females crossed the tunnel at higher frequency and did not prefer a side to lay their eggs (Figure E), suggesting that mated females actively search others when isolated and are attracted to lay their eggs with others despite the resulting competition. These results therefore suggest that mated females actively search others when isolated and are attracted to lay their eggs with others despite the resulting competition.

### Light interacts with social context to modulate the timing of egg-laying

Egg-laying is regulated by the circadian clock, which is synchronized by environmental cues such as light (Manjunatha et al., 2008). Egg-laying in *Drosophila* occurs mainly at night (Howlader and Sharma, 2006; Manjunatha et al., 2008; Sheeba et al., 2001), which might act to protect eggs against UV damage and desiccation (Zhu et al., 2014), as well as predation from parasitic wasps (Howlader and Sharma, 2006). We noticed that grouped females were not only faster in initiating egg-laying than lone females, but also started laying eggs during the day while lone females waited for the onset of darkness. Social context and light might thus interact to determine timing of egg-laying. To test this hypothesis, we quantified the number of eggs laid in bins of two hours over 48 hours in 12:12 Light:Dark (LD) conditions, both for single and grouped females. When the experiment started at Circadian Time (CT) 5, the focal females in a group started laying eggs at CT7, which is during the light phase, and slowed down during the dark phase to resume again during the next light phase, and then laid eggs at a steady rate through the second dark phase (Figure 2A). By contrast, isolated females started laying eggs at CT12, which is when darkness began. Isolated females laid eggs at an accelerated rate in the first part of the dark phase, catching up with grouped females halfway through the dark phase after which they stopped laying eggs until the beginning of the next dark phase (Figure 2A). This egg-laying timing shift was also repeated during the second light/dark cycle. Although isolated females started laying eggs later than grouped females, both isolated and grouped females laid approximately the same total number of eggs after 24h and 48h (Supp. Figure S2A). Results show that solitary egg-laying coincides with the dark phase and suggest that isolated females retain their eggs until darkness. Grouped females however lay eggs already during the light phase, which leads to depressed egg-laying during the first dark phase. That grouped females then lay at a steady rate through the next light and dark phases suggests that grouped females are insensitive to the effect of light on egg-laying behaviour. These results support the hypothesis that light conditions and social context interact to determine the timing of egg-laying.

**Figure 2:**
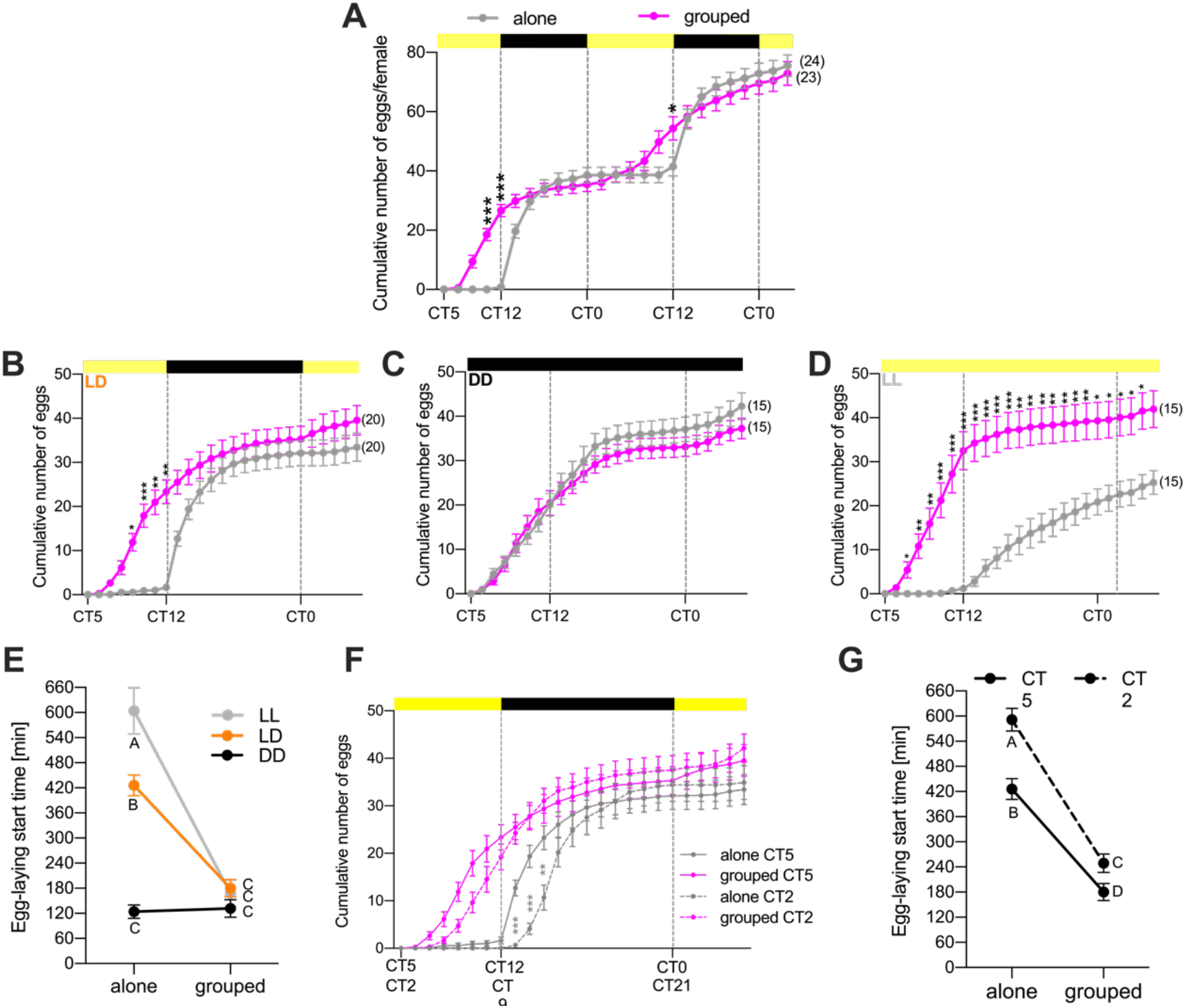
Light interacts with social context to modulate egg-laying. **(A)** Cumulative number of eggs in 48h of a wild-type *Oregon-R* mated female alone or in groups with 5 *Oregon-R* males. Cumulative number of eggs was counted every 2 hours during 48 hours on a 12h/12h light/dark cycle. Replicates per group: 23-24. **(B-D)** Cumulative number of eggs in 24h of a wild-type *Oregon-R* mated female alone or in groups with 5 *Oregon-R* males. Cumulative number of eggs was counted every hour during 24 hours in different light/dark conditions: **B-** 12h/12h light/dark cycle, **C-** 24h dark, **D-** 24h light. Replicates per group: 15-20. **(E)** Onset of egg-laying of a wild-type *Oregon-R* mated female alone or grouped in different light/dark conditions. LD: 12h/12h light/dark, DD: 24h dark, LL: 24h light. Replicates per group: 15. **(F)** Cumulative number of eggs in 24h of a wild-type *Oregon-R* mated female alone or in groups with 5 *Oregon-R* males. Cumulative number of eggs was counted every hour during 24 hours in 12h/12h light/dark conditions. The experiment was set up at CT2 or CT5 with a dark phase starting respectively at CT9 or CT12. Replicates per group: 20. **(G)** Onset of egg-laying of a wild-type *Oregon-R* mated female alone or grouped in different experimental onset time: beginning of the experiment at CT2 or CT5. Replicates per group: 19-20. Error bars indicate standard error of the mean. Different letters or stars indicate differences between alone vs grouped and between CT2 and CT5 (*** p<0.001, ** p<0.01, * p<0.05). For full statistical analysis see **Supplementary Table S1.**

To directly test the hypothesis that light and social context interact to determine the onset of egg-laying, we set up single or grouped flies in constant light or dark conditions. In dark conditions, single females laid eggs as fast as grouped females and had the same egg-laying dynamics when in constant dark conditions (Figure 2C) showing that single females are physiologically able to lay eggs as fast as grouped females, but that they delay egg-laying during day-time. In constant light conditions, isolated females started laying eggs after 8 hours of exposure to constant light (Figure 2D) and laid fewer eggs than grouped females after 24h (Supp. Figure S2B), showing that light reduces female fecundity when alone but that light cannot fully inhibit egg-laying. Further analysis confirmed a strong interaction of social context and light (*p<0.001*) in egg-laying start-time (Figure 2E). These data show that the onset of egg-laying is inhibited by light, but that this inhibition is lifted by the presence of other females. Social context and light thus interact to modulate egg-laying start time.

Is the effect of light direct or is it mediated by its entrainment of the circadian clock? To begin to investigate the relative action of the circadian clock vs light on social-context-dependent egg-laying dynamics, we introduced a 3h-shift by changing both the start-time of the experiment from CT5 to CT2 and the onset of the dark phase from CT12 of CT9 (Figure 2F). In these phase-shifted conditions, grouped females started laying eggs in the light phase while light inhibited egg-laying in isolated females who waited for darkness to lay their eggs in both experimental timings CT2 and CT5. However, CT2 females from both single and grouped conditions shifted egg-laying dynamics by 2 hours compared to CT5 females. CT2 isolated females did not lay eggs when light was off and waited 2 hours in the dark before starting egg-laying unlike CT5 isolated females (Figure 2F&G). Egg-laying latency comparisons between CT2 and CT5 experimental times indicated that the start-time of egg-laying of isolated and grouped females is significantly delayed by 1.5h when the experiment begins at CT2 (Figure 2G). These results show that the circadian clock influences egg-laying dynamics of *Drosophila* females whether alone or in a group. Further analysis revealed a weak interaction of social context and circadian time (*p = 0.044*) in egg-laying start-time. Taken together, our data suggest that the circadian clock plays a modest role in modulating egg-laying start-time in both grouped and single females. Light itself inhibits egg-laying start-time only in isolated females, grouped females being insensitive to light conditions. Social context can thus trump environmental conditions to modulate *Drosophila* female egg-laying.

### Females rely on visual cues to modulate timing of egg-laying

To understand the mechanisms by which females adjust the onset of egg-laying to the social environment, we started by investigating which cues and sensory inputs regulate this behaviour. We first explored the specificity of the modulation of egg-laying by testing whether *D. melanogaster* individuals from different sex, mating status and species can trigger an advancement in egg-laying behaviour of a focal female. To control for males inducing an egg-laying delay by harassing the females, we used both wild-type males and *fruitless (fru)* mutant males, who lack courtship behaviour (Demir and Dickson, 2005). Females advanced egg-laying in the presence of males regardless of their courtship behaviour (Figure 3A). Females also advanced egg-laying in the presence of virgin females (Figure 3A). To test whether other *Drosophila* species were able to trigger an egg-laying advancement, we tested females either alone or grouped with 5 males from different *Drosophila* species. *Drosophila* species included both close and evolutionary distant ones, as well as species ranging in size from 1.5 to 4 mm and in colour from beige to dark (Chyb and Gompel, 2013). Focal *D. melanogaster* females advanced the timing of their first eggs when grouped with any of the *Drosophila* species tested compared to when alone (Figure 3B). Because cuticular hydrocarbons (CHCs) vary according to the sex, mating status and *Drosophila* species (Billeter and Levine, 2015; Billeter and Wolfner, 2018; Everaerts et al., 2010; Ferveur, 1997), these findings suggest recognition of species, sex and mating status is not required for egg-laying advancement. To verify this, we assayed egg-laying behaviour of a focal wild-type female in the presence of oenocyteless flies (oe^-^), which lack CHCs but otherwise behave like wild-type animals (Billeter et al., 2009). Mated females did advance egg-laying when grouped with both control and oe^-^ flies from both sexes (Figure 3C). CHCs are thus not necessary for egg-laying advancement. In addition, overall fecundity is not modulated by group composition, as grouped females laid approximately the same number of eggs in 24h regardless the sex, the mating status and the species of group members (Supp. Figure S3). Egg-laying advancement is thus not triggered by sex, colour, or species recognition cues. This indicates that a more general cue informs this behaviour.

**Figure 3:**
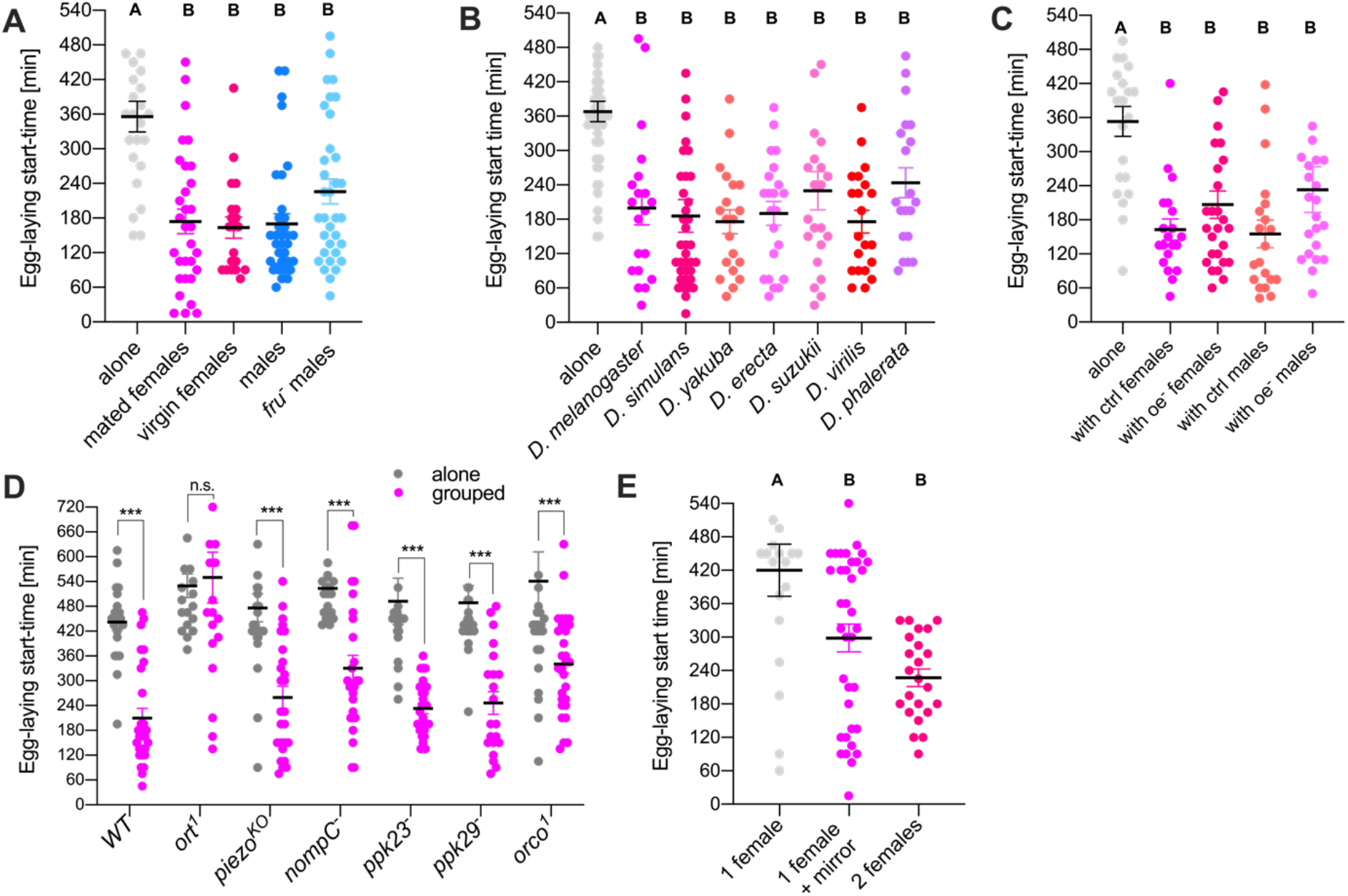
Group density is the cue that modulates egg-laying and is detected by vision. **(A)** Onset of egg-laying of a wild-type *Oregon-R* mated female alone, grouped with 5 mated females, with 5 virgin females, with 5 males or with 5 *fruitless* males. Replicates per group: 20-34. **(B)** Onset of egg-laying of a wild-type *Oregon-R* mated female alone or grouped with 5 males from different *Drosophila* species. Species were from different relatedness and were ranged from the least distant to the most distant related one from *D. melanogaster*: *D. simulans, D. yakuba, D.erecta, D. suzukii, D. virilis, D. phalerata*. Replicates per group: 20-37. **(C)** Onset of egg-laying of a wild-type *Oregon-R* mated female alone, grouped with 5 oenocyteless mated females, 5 oenocyte-less males, control females or control males. Replicates per group: 20-27. **(D)** Onset of egg-laying of a mated female alone or grouped with 5 males. Mated females were wild-type *Oregon-R* or mutant for different sensory modalities: *ort*^*1*^ (vision), *piezo*^*KO*^ and *nompC* (touch), *ppk23*^*-*^ and *ppk29*^*-*^ (taste), *orco*^*1*^ and *or47b*^*3*^ (smell). To remove variation arising from genetic background, mutant lines were all placed into an *Oregon-R* genetic background by 10 generations of backcrossing to an *Oregon-R* strain. Replicates per group: 20-29. **(E)** Onset of egg-laying of a wild-type *Oregon-R* mated female grouped with another *Oregon-R* mated female or alone with or without a circular mirror applied along the inside of the egg-laying dish. Replicates per group: 19-36. Error bars indicate standard error of the mean. Different letters or stars indicate differences between alone vs grouped (*** p<0.001, ** p<0.01, * p<0.05). For full statistical analysis see **Supplementary Table S1.**

To indirectly uncover the cue(s) that regulate(s) egg-laying advancement, we tested the necessity of a series of sensory modalities including vision, touch, taste and smell for this behaviour by testing egg-laying latency of mutant females alone or in a group (Fig 3D). Two mutants for touch were tested: *Piezo*^*KO*^, a genetic knockout of the receptor detecting strong touch (Coste et al., 2012; Kim et al., 2012) and *nompC*, a mutant with altered mechanosensory transduction channels (Walker, 2000). For olfaction, *Orco*^*1*^ mutants lacking the function of classical odorant receptors (Larsson et al., 2004; Silbering et al., 2011) were tested. For taste, *Ppk23*^*-*^ and *ppk29*^-^ mutants that are necessary for CHC sensing were tested to further assay the necessity of pheromone sensing (Lu et al., 2012; Thistle et al., 2012; Toda et al., 2012). None of these sensory modality mutants showed altered start-time of egg-laying revealing that touch, smell and taste are not necessary for social context-dependent egg-laying modulation (Figure 3D). Female *ort*^*1*^ mutants, which lack the first neurotransmission from photoreceptors and are thus blind (Gengs et al., 2002) did not advance egg-laying compared to isolated females (Figure 3D). This suggests that vision is the sensory modality necessary to sense social context. This was further shown by placing a wild-type mated female alone in a dish surrounded by a flexible circular mirror reflecting her image or in group with another female. Females advanced egg-laying when grouped with another fly but also when alone surrounded by a mirror (Figure 3E). These results showed that, even though the response is not as strong as the presence of another fly, her own reflection in the mirror was sufficient to mimic the presence of a fly and to trigger a fast oviposition, indicating that visual input is sufficient for group detection and for egg-laying advancement.

### The motion detection circuit is necessary for egg-laying advancement

Having established that vision is necessary and sufficient for egg-laying advancement, we next determined whether females modulate egg-laying start-time depending on group size or density. To do that, we tested a focal female either alone or grouped with 5 females on a food patch placed in different size arenas. Start-time of egg-laying for females alone did not change with increasing dish sizes showing that area size itself does not impact egg-laying timing (Figure 4A). However, timing of egg-laying of a focal female in a group of 6 females increased with arena size ultimately reaching an egg-laying time similar to that of an isolated female (Figure 4A). We interpret these results as meaning that egg-laying timing is density -not group-size-dependent. Further correlation analysis did not show a linear relationship between egg-laying timing of grouped females and density but showed a decreasing exponential relationship (Figure 4B), suggesting an active regulation of egg-laying by density. To gain more insight in how females assess density in these groups, we used automated behavioural tracking to determine the position of individual females in different size arenas and quantify their social interactions. As group density decreased, the number of physical contacts decreased (Figure 4C) and inter-individual distance increased (Figure 4D), both in a linear fashion. This linear increase in contact, taken together with the observation that contact sensing is not necessary for modulation of egg-laying (Figure 4D), suggest that density-which affects egg-laying in an exponential manner-is not sensed by contact.

**Figure 4:**
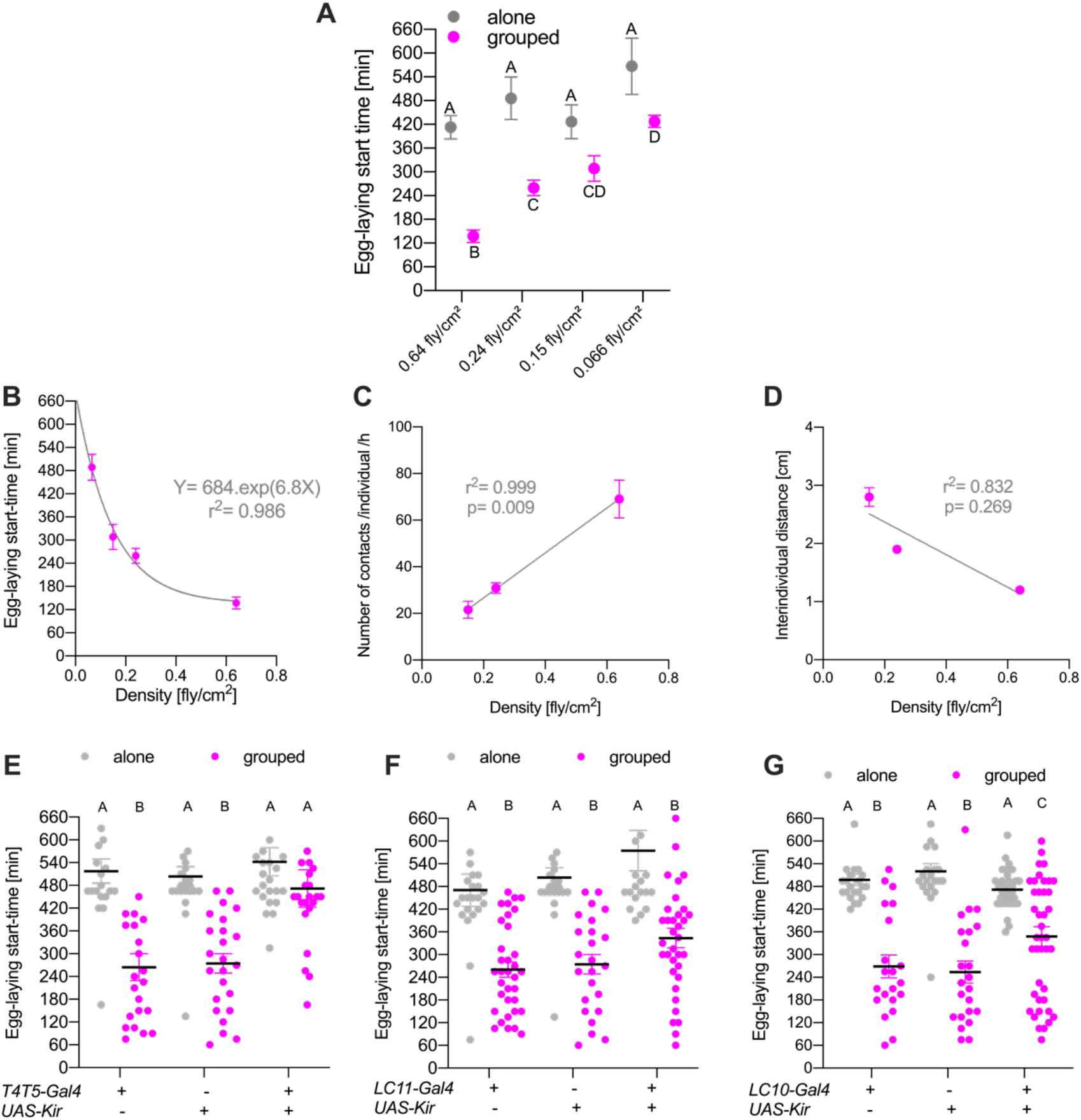
Females detect group density via T4T5 and LC10 size and motion sensitive neurons. **(A)** Onset of egg-laying of a wild-type *Oregon-R* mated female alone or in group with 5 other *Oregon-R* mated females in different petri dishes. Different sizes of petri dishes (30×10mm, 55×14mm, 90×14mm, 40×20.6mm) were used for the experiment but the same food patch size was used in all conditions. Replicates per group: 14-23. **(B)** Average onset of egg-laying of a wild-type *Oregon-R* mated female in group with 5 other *Oregon-R* mated females in different densities induced by different sizes of petri dishes (30×10mm, 55×14mm, 90×14mm, 40×20.6mm). Correlation between start-time of egg-laying and density was also established. **(C)** Average number of contacts of a wild-type *Oregon-R* mated female in a group with 5 other *Oregon-R* mated females in different densities induced by different sizes of petri dishes (30×10mm, 55×14mm, 90×14mm, 40×20.6mm). Correlation between number of contacts per individual and density was also established. **(D)** Average inter-individual distance between 6 grouped wild-type *Oregon-R* mated females in different densities induced by different sizes of petri dishes (30×10mm, 55×14mm, 90×14mm, 40×20.6mm). Correlation between inter-individual distance and density was also established. **(E)** Onset of egg-laying of a mated female alone or grouped with 5 males. Mated females with silenced T4T5 neurons (*UAS-Kir x T4T5-gal4*) or functional T4T5 neurons (controls: *UAS-Kir x OR* and *T4T5-gal4 x OR*) were tested. Replicates per group: 20-24. **(F)** Onset of egg-laying of a mated female alone or grouped with 5 males. Mated females with silenced LC11 neurons (*UAS-Kir x LC11-gal4*) or functional LC11 neurons (controls: *UAS-Kir x OR* and *LC11-gal4 x OR*) were tested. Replicates per group: 20-35. **(G)** Onset of egg-laying of a mated female alone or grouped with 5 males. Mated females with silenced LC10 neurons (*UAS-Kir x AD-DBD LC10-gal4*) or functional LC10 neurons (controls: *UAS-Kir x OR* and *AD-DBD LC10-gal4 x OR*) were tested. Replicates per group: 21-42. To remove variation arising from genetic background, *T4T5-gal4, LC11-gal4, AD-DBD LC10-gal4, UAS-Kir* lines were all placed into an *Oregon-R* genetic background by 10 generations of backcrossing to an *Oregon-R* strain. Error bars indicate standard error of the mean. Different letters indicate differences between alone vs grouped. For full statistical analysis see **Supplementary Table S1.**

Our sensory mutant data suggest that females detect group density via vision leading to egg-laying advancement. As flies trigger earlier egg-laying in focal females independent of fly size and coloration, we hypothesise that movement is the cue that is detected. To establish whether movement is a social cue for egg-laying advancement, we tested mutants for the first stage of computation of visual motion, the columnar cells T4 and T5 neurons (Borst et al., 2020). T4 and T5 neurons exist in four subtypes, each corresponding to motion in one of the four cardinal directions and project their axons in the different layers of the lobula plate which contain wide-field motion-sensitive neurons (Borst et al., 2020; Maisak et al., 2013; Mauss et al., 2017; Schnell et al., 2012). To test our hypothesis, we performed an egg-laying experiment and silenced T4 and T5 cells by expressing Kir2.1 in mated females with the presence or absence of a group. Unlike the UAS and gal4 controls, females did not advance egg-laying in a group when T4 and T5 were blocked (Figure 4E), showing that T4 and T5 are necessary for group sensing. Lobula columnar (LC) neurons are an entry point for the circuit-level study of visual responses since it is the first layer where visual information about objects is decoded (Ferreira and Moita, 2020; Ribeiro et al., 2018; Wu et al., 2016). Having identified motion cues of group members as the leading source of the group effect on egg-laying, we investigated the role of visual projection neurons responsive to the movement of small objects like a flies: LC10 and LC11 (Ferreira and Moita, 2020; Keleş and Frye, 2017; Ribeiro et al., 2018; Wu et al., 2016). LC neurons were silenced in females by expressing Kir2.1 and by testing the focal female for egg-laying latency alone or in groups with wild-type males. As UAS- and gal4-controls, silenced-LC11 females laid eggs faster in group than when alone, indicating that LC11 neurons are not involved in group detection leading to egg-laying advancement (Figure 4E). When silencing LC10 neurons in mated females, we still found a decreased egg-laying start-time when females are in group than when alone (Figure 4G).

However, egg-laying advancement of grouped females is not as strong as of the two controls (i.e. UAS and gal4 controls), the majority of *UAS-Kir x LC10gal4* grouped females waiting for darkness to start laying eggs. In other words, silencing LC10 neurons rendered females less sensitive to the motion of others, revealing LC10 neurons as part of the visual neuronal circuitry for group density detection. The variation found in silenced-LC10 females can either be linked with incomplete penetrance of Kir2.1 expression or the meaning that LC10 are partially involved in group detection and that other neurons might be also involved in sensing group in the context of egg-laying advancement. Taken together, these experiments reveal that females detect social context through a motion detection pathway which modulates their egg-laying timing, and that T4/T5 neurons and upstream LC10 neurons are involved in the neuronal circuit that detects other flies motion leading to a female egg-laying modulation.

### Social context modulates oogenesis and ovulation

We showed that while single females delay egg-laying until darkness, grouped females lay eggs during the day, resulting in faster egg-laying in a group than alone. As egg production can be regulated both at the oogenesis and ovulation level, differences in egg-laying timing in single versus grouped females could thus be explained by the lifting of an inhibitory effect of light on either of these two processes. To investigate this, recently mated females were placed alone or in a group and their ovaries and oviducts were dissected to quantify oogenesis and ovulation at two time points after mating: 2 hours and 6 hours. These two time points are respectively the time at which grouped females begin to lay eggs, and 1 hour before darkness when isolated females start laying eggs under our standard assay conditions. Females were also placed and dissected in full dark conditions 2 hours after mating - the time at which both isolated and grouped females start laying eggs. At the first time point, 2 hours post-mating, we found a much higher probability of ovulation in grouped than in isolated females (Figure 5A), indicating that social context modulates ovulation. Similarly, one hour before darkness (6 hours post-mating), we still found a decrease in ovulation rate of isolated females compared to the grouped ones (Figure 5A), suggesting that light inhibits ovulation in alone conditions. To test the hypothesis that light inhibits egg production specifically in isolated females, oviducts of lone and grouped females were dissected under full dark conditions 2 hours after mating. Results showed that the percentage of ovulation is the same for both isolated and grouped females, supporting the hypothesis that light delays ovulation in isolated females as opposed to speeding it up in grouped females (Figure 5A).

**Figure 5:**
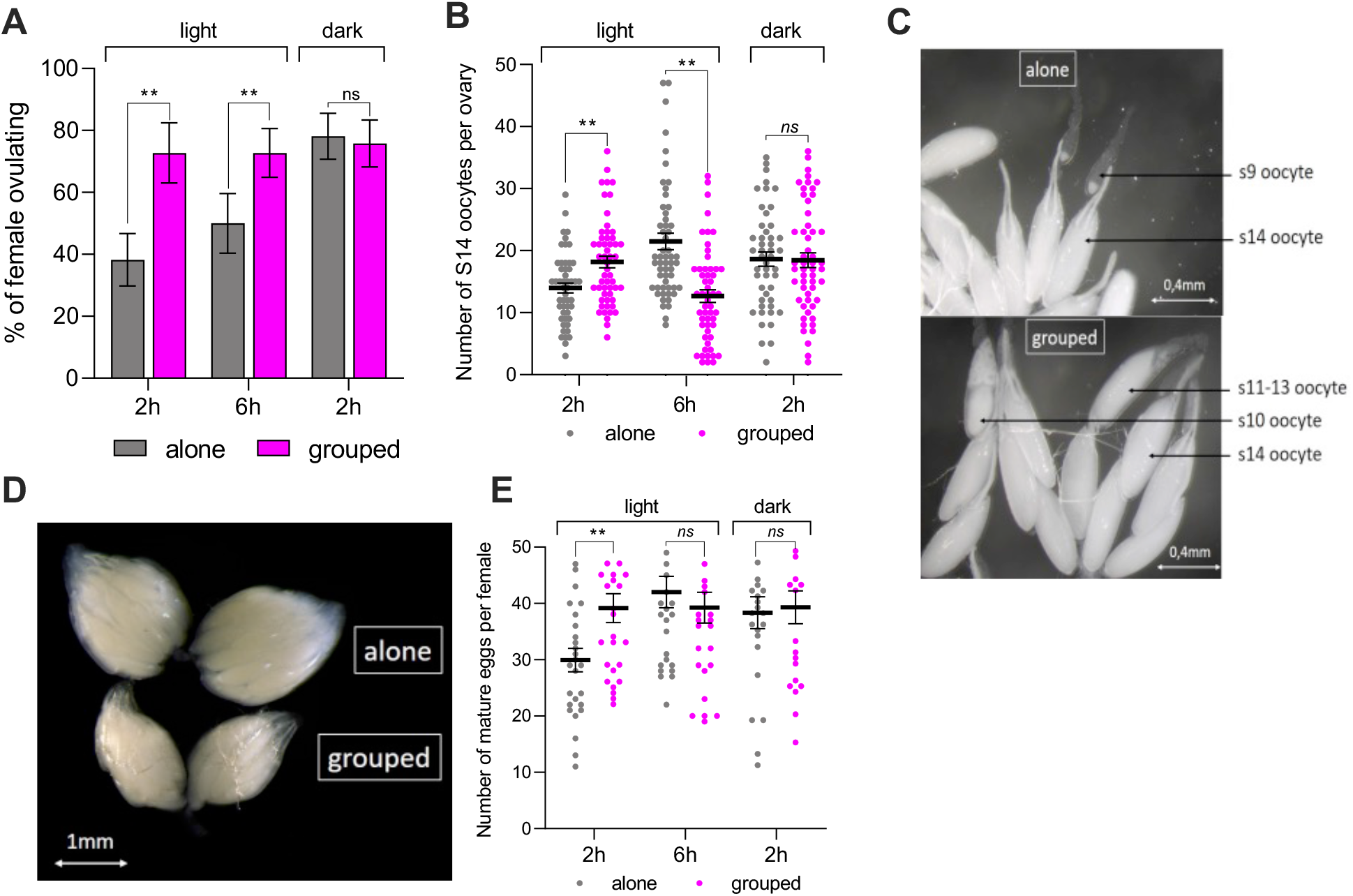
Light inhibits oogenesis and egg production in isolated females while the presence of a group lifts the inhibition leading to oogenesis, ovulation and oviposition advancements. **(A)** Ovulation rate of isolated and grouped wild-type *Oregon-R* females 2h and 6h post-mating under light and dark conditions. Percentage of females that are ovulating was measured by monitoring the presence or absence of an egg in the oviducts. Replicates per group: 22-34. **(B)** Number of stage 14 oocytes per ovary of isolated or grouped wild-type *Oregon-R* females 2h and 6h post-mating under light and dark conditions. Stage 14 is the stage where oocytes are mature and ready to be laid. 48 to 54 ovaries were dissected corresponding to 24 to 27 females. Replicates per group: 48-56 ovaries. **(C)** Photos of ovarioles from an isolated (above) and a grouped (below) female’s ovary 2h post-mating under light condition. Grouped ovarioles contained more s14 oocytes than isolated ovarioles. **(D)** Photo of ovaries from an isolated (above) and a grouped (below) female 6h post-mating under light condition. Isolated ovaries are bigger than the grouped ones. **(E)** Total number of mature eggs per female of isolated and grouped wild-type *Oregon-R* females 2h and 6h post-mating under light and dark conditions. To quantify the total number of mature eggs per female, the number of eggs laid after mating and before dissection, the number of eggs in oviduct and the number of eggs at stage 14 in ovaries were counted. Replicates per group: 22-27. Error bars indicate standard error of the mean. Different letters or stars indicate differences between alone vs grouped (*** p<0.001, ** p<0.01, * p<0.05). For full statistical analysis see **Supplementary Table S1.**

We then looked at mated female’s oogenesis under light conditions depending on social context. Two hours post-mating, ovaries of grouped females contained more stage 14 oocytes than those of isolated females, showing that being in a group stimulates female oogenesis under light conditions (Figures 5B-C). At 6 hours post-mating, isolated females had more mature stage 14 oocytes in ovarioles than grouped females (Figure 5B). This increased number of stage 14 oocytes in isolated females, resulting in visibly bigger ovaries (Figure 5D), is explained by the accumulation of mature oocytes in the ovarioles, consistent with the interpretation that ovulation is inhibited when females are alone in the light. The reduced number of stage 14 oocytes in grouped females is explained by the fact that grouped females are in the process of egg-laying and thus ovulating these mature oocytes. Oogenesis did not differ between isolated and grouped females in dark conditions (Figure 5B), backing the hypothesis that isolated females delay egg production under light conditions. These results also suggest that, 6 hours after mating, females alone caught up egg maturation and production and accumulated mature stage 14 eggs in ovaries, waiting for darkness to start laying eggs.

To check whether isolated females are able to produce as many mature eggs as grouped females in 6 hours –despite keeping them in their ovaries-, we counted the number of eggs laid after mating and before dissection, the number of eggs in oviduct and the number of eggs at stage 14 in ovaries. 6 hours after mating, isolated and grouped females produced the same number of mature eggs (Figure 5E), confirming that social environment does not modulate overall fecundity. Isolated and grouped females both produced about 40 mature eggs, which look to be the maximum mature eggs able to be produced by a female in this period of time. However, 2 hours after mating, grouped females already reached this potential maximal number of mature eggs produced by a female (Figure 5E), highlighting that social environment modulates the rate of egg production. Quantification of mature eggs 2 hours post-mating under full dark conditions confirmed that isolated females are able to produce the maximal number of eggs as fast as grouped females when in darkness (Figure 5E).

Taken together our data show that light inhibits egg production, but this inhibition can be lifted by the presence of other females. Isolated females delay egg-laying retaining eggs in ovaries when light, while grouped females advance oogenesis, ovulation and thus oviposition.

### Juvenile hormone modulates oogenesis in response to social context and light

Juvenile hormone (JH) is central to the regulation of oogenesis in insects and in Droosphila, making it a candidate for the social modulation of oogenesis (Meiselman et al., 2018; Soller et al., 1999; Terashima et al., 2005; Terashima and Bownes, 2004). We hypothesised that light reduces JH level but that being in a group increases JH levels, which allows females to lay eggs during the light phase when in group. To test whether JH rescue is sufficient to salvage reproductive output of isolated females under light conditions, we applied the JH analogue Methoprene in isolated females immediately after mating. Ovaries and oviducts of isolated mated females in light conditions were dissected 2h post-mating and Methoprene application. Addition of Methoprene in isolated mated females resulted in the production of more stage 14 oocytes than isolated mated females supplemented with solvent control. The Methoprene-treated females produced a similar amount of s14 eggs as grouped females (Figure 6A). That application of JH analogue is sufficient to stimulate oogenesis in single females despite inhibition coming from the light suggests that increased JH is the endogenous mechanism that modulates the effect of social context on oogenesis. This suggestion needs to be backed up by *in vivo* measurement of JH titer in single vs grouped females; however, our current attempt at quantifying JH failed to reveal a statistically significant effect, probably due to small effect sizes. In addition, despite the effect on oogenesis, Methoprene-treated single females had neither higher ovulation rates (Figure 6B) nor laid eggs as fast as grouped females (Figure 6C), showing that JH is not the only hormone that modulates the effect of social context on egg-production.

**Figure 6:**
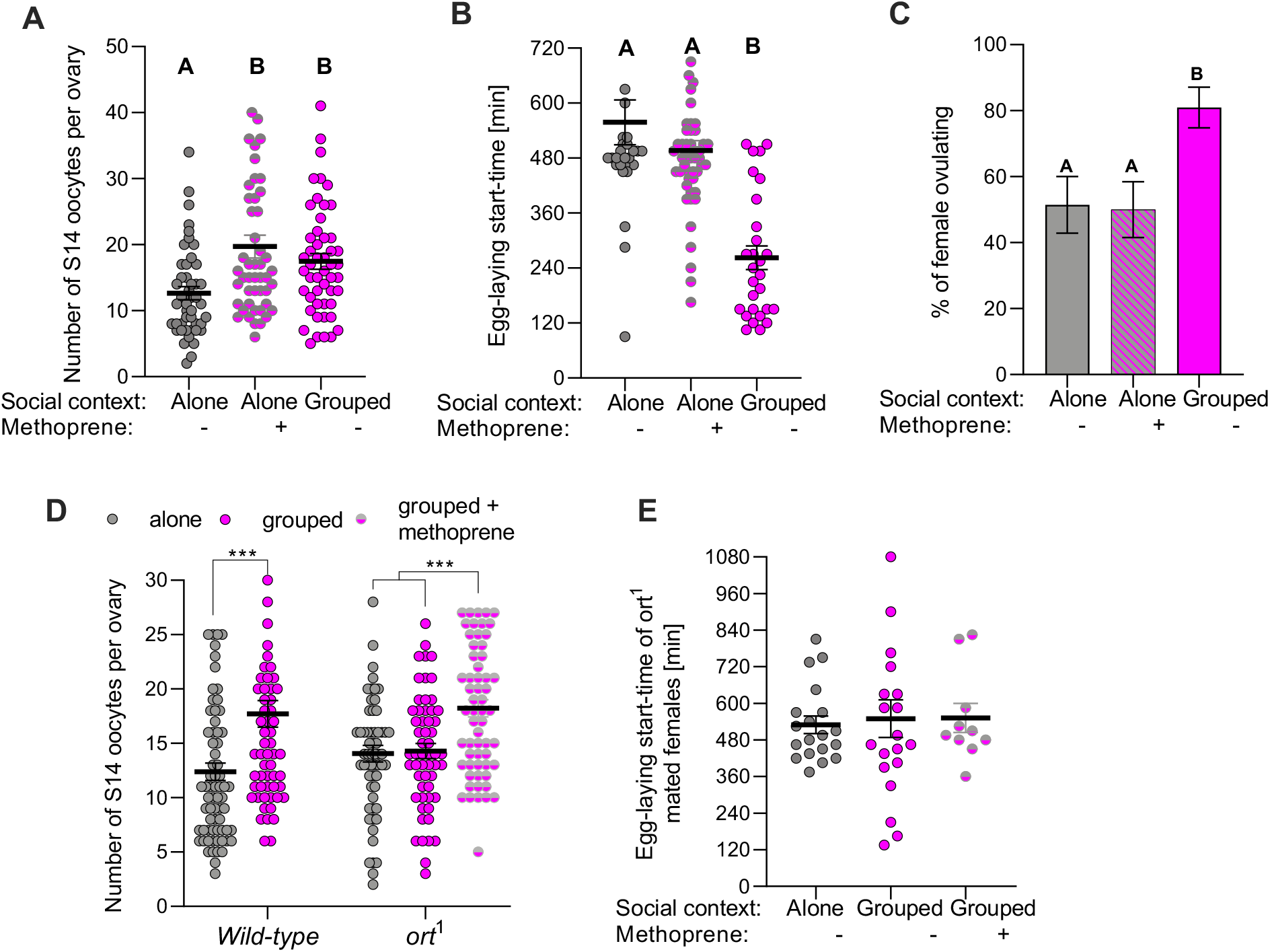
Juvenile hormone production stimulates female oogenesis in groups, lifting the inhibition of light. **(A)** Number of oocytes at stage 14 per ovary of a wild-type *Oregon-R* mated female alone or grouped with 5 males, 2h after application of methoprene or acetone under light condition. Replicates per group: 46-50. **(B)** Onset of egg-laying of a wild-type *Oregon-R* mated female alone or grouped with 5 males, 2h after application of methoprene or acetone under light condition. Replicates per group: 28-49. **(C)** Ovulation rate of a wild-type *Oregon-R* mated female alone or grouped with 5 males, 2h after application of methoprene or acetone under light condition. Replicates per group: 35-42. **(D)** Number of oocytes at stage 14 per ovary of a wild-type *Oregon-R* and a blind ort^1^ mated female alone or grouped with 5 males, 2h post-mating under light condition, with or without application of methoprene. Replicates per group: 60-70. **(E)** Onset of egg-laying of a mated female alone or grouped with 5 males, 2h post-mating under light condition, with or without application of methoprene. Mated females were wild-type *Oregon-R* or mutants for vision (*ort*^*1*^). Replicates per group: 10-20. To remove variation arising from genetic background, *ort*^*1*^ line was placed into an *Oregon-R* genetic background by 10 generations of backcrossing to an *Oregon-R* strain. Error bars indicate standard error of the mean. Different letters or stars indicate differences between alone vs grouped (*** p<0.001, ** p<0.01, * p<0.05). For full statistical analysis see **Supplementary Table S1.**

Taken together our data demonstrate that *Drosophila* integrates light cues that normally inhibit egg-laying with social cues detected by vision - in particular the motion pathway- leading to changes in hormonal titer that ultimately regulate oogenesis. To directly test this, we applied Methoprene on *ort*^*1*^ mutant blind females to restore oogenesis in these females. As shown previously, *ort*^*1*^ females do not advance egg-laying when in a group compared to wild-type females (Figure 3D). Quantification of oogenesis in these single and grouped blind females revealed a reduced number of stage 14 oocytes compared to wild-type females when grouped (Figure 6D). Methoprene-treated grouped blind females produced significantly more stage 14 oocytes than the solvent control ones and produced a similar amount of s14 eggs as wild-type grouped females (Figure 6D), showing that JH analog application stimulated oogenesis and restored oogenesis in grouped blind females. As expected, grouped blind females did not lay eggs faster in a group after methoprene treatment (Figure 6E), confirming our finding that JH analog application stimulates and restores oogenesis but does not advance egg-laying. Further studies will be needed to reveal what other hormones/neurons /pathways are working in combination with JH to promote ovulation and egg-laying advancement.

## Discussion

In this study, we investigated the mechanisms through which fundamental behavioural and physiological aspects of female reproduction are modulated by the presence of others in *D. melanogaster.* We demonstrated that grouped females start laying eggs during the light phase of the day, while isolated females solely lay them during the night. The presence of flies from any sex, mating status or species can trigger these responses. We also showed that start-time of egg-laying is density-dependent and that females detect the presence of others via vision through the motion detection pathway. Finally, we found that being in a group lifts the inhibitory effect of light resulting in grouped females laying eggs during the day, while isolated females retain their eggs until darkness. Our study further revealed that this egg-laying modulation by group is connected to a lifting of the inhibition of light on oogenesis by stimulating a hormonal pathway involving juvenile hormone.

### Communal egg-laying in *Drosophila melanogaster*: a case of competition or cooperation?

*D. melanogaster* females lay eggs faster in a group than alone. That this behaviour leads to competition among offspring is revealed by decreased egg-to-adult survival, especially for eggs that arrive later than others on the substrate (Figure 1D). This suggests that grouped females lay eggs fast to give a competitive advantage to their offspring, who can first exploit the finite resources. We found several offspring that were still at the larval developmental stage 15 days after egg-laying (whereas without competition, complete larval and pupal development normally takes 10 days under these conditions). This suggests larvae could not all obtain enough nutrients for completing development. By taking up the nutrients that are necessary for their development early, larvae have a higher chance of survival when resources are limited, poor or quickly decaying, as is the case for the food substrates exploited by *Drosophila* in nature. Our findings are in line with the documentation that, at high density, individuals compete for food, even leading to cannibalism between larvae (Narasimha et al., 2019; Vijendravarma et al., 2013b), causing a decrease of offspring survival (Allee, 1927b; Courchamp et al., 1999; Etienne et al., 2002b; Stephens and Sutherland, 1999; Wertheim et al., 2002c). In our experiments, we noticed (but did not quantify) a majority of larvae eating each other when dishes were composed of hosts and guests that were out of synchrony. Lack of nutrients and cannibalism could thus explain the very low survival of guest offspring when they were placed on the poor substrate later than hosts.

That our assays revealed competition leading to lower survival is paradoxical given that flies actively attract others for communal egg-laying (Billeter and Levine, 2015; Billeter and Wolfner, 2018; Brown et al., 2003; Dawson et al., 2018; Duménil et al., 2016b; Koto et al., 2015; Lihoreau et al., 2009; Wertheim et al., 2002a; White et al., 2017). We complement this knowledge by showing here that females actively search others and are attracted to lay their eggs with others despite the resulting competition among their offspring (Figure 1E). Larval aggregation is a behaviour that results in a higher offspring survival thanks to cooperation in fending off fungal growth (Trienens et al., 2017; Wertheim et al., 2002c). There is therefore a trade-off between finding a group to increase chance of offspring survival via cooperation and finding a group that is not too big to avoid competition and a dramatic decrease of survival. It is likely that the absence of competitors (e.g. fungi) in our laboratory assays has masked the beneficial effects of communal egg-laying. We thus predict that flies in the wild experience benefits of communal egg-laying, yet need to hurry egg-production to avoid competition. In this scenario, faster egg-laying might be seen as a mechanism of competition avoidance. Additionally, it also achieves greater synchronization of egg-laying between females, increasing larval survival.

Because the benefit of social interactions depends on the identity of group members, social interactions should be instructed by recognition of cues indicating sex, mating status or species. Many organisms use chemical signals to recognize others (Wyatt, 2014), including *D. melanogaster*, which use CHCs signals to distinguish conspecific individuals from others (Ferveur, 2005; Jallon, 1984; Seeholzer et al., 2018; Wyatt, 2014). CHCs mediate chemical communication for both sex and species recognition and instruct courtship, mating interactions and aggregation behaviours (Billeter et al., 2009; Billeter and Levine, 2013; Duménil et al., 2016b; Ferveur, 2005; Hall, 1994; Jallon, 1984; Scott, 1994; Wicker-Thomas, 2007). Here we surprisingly demonstrated that the presence of flies from any sex, mating status or species can trigger a fast oviposition in females, showing that detailed recognition is not required for egg-laying advancement. All tested flies, even from species including both close and evolutionary distant ones, as well as different sized and colours, triggered this egg-laying behavioural response. In German cockroaches, interactions with other species (e.g. katydid, brown-banded cockroach, or camel cricket) of similar or larger size have also been found to stimulate oocyte development and female reproduction (Uzsák and Schal, 2013). In the wild, multiple *Drosophila* species can coexist on the same fruits (Sevenster, 1996; Symonds and Wertheim, 2005; Wertheim et al., 2006). Larvae from all these species can both potentially compete for the same resources and cooperate in preventing the growth of harmful microorganisms. Precise recognition might thus not be required.

The presence of males also leads to an egg-laying advancement. In mammals, the presence of males modifies female reproduction by accelerating puberty and reproduction in several species of rodents, farm animals and in a primate (Drickamer, 1983; Muller-Schwarze, 1984). This behaviour can be explained by the fact that the faster females are ready to be fertilized, the more likely they will find an available male. In German cockroach females, being with females or males result in the stimulation of oocyte development (Gadot et al., 1989), as we reveal here in *Drosophila*. Finding that the presence of males trigger egg-laying advancement in *Drosophila* females can be explained by the fact that, in addition of being sexual partners, the presence of males on the substrate may bring yeasts and feces which may respectively increase the nutritional value of a food patch and hamper the growth of noxious microorganisms (Wertheim et al., 2002c).

### The motion pathway as a modality to detect group density

We show that females modulate egg-laying timing according to group density. The lower the density, the slower egg-laying timing. A lower density implies that females encounter others less frequently, decreasing their interaction rates and modifying social network properties (Rooke et al., 2020). Our tracking analysis indeed revealed that the number of contacts and social distance changed with density, making the focal female interacting less with others as density decreased (Figure 4 A-D). Vision is the main sensory modality necessary to sense the social environment in the context of egg-laying advancement. Vision is essential for other *Drosophila* social behaviours. It determines whether to approach a moving object and initiate courtship by allowing discrimination of objects that have the dimension of a fly (Agrawal et al., 2014; Bidaye et al., 2020; Ribeiro et al., 2018). Flies also detect group presence via vision and gradually respond to group size by displaying lower freezing responses as group size increases (Ferreira and Moita, 2020). Moreover, flies detect the presence of parasitoid wasps through visual cues, which lead to egg-retention and ovarian apoptosis (Kacsoh et al., 2015). Vision thus seems to be an essential modality by which *Drosophila* females sense their environment and regulate egg-laying behaviour: blocking eggs in the presence of parasites and stimulating egg-laying in the presence of flies. Despite our finding that most flies can trigger egg-laying advancement, we expect flies to be able to discriminate conspecific against predators such as parasitoid wasps. Further studies to dissect which visual pathway differentiates flies vs parasitoid wasps are now possible by testing egg-laying advancement vs retention in response to different tyopes of visual cues.

Females detect social context specifically through the motion detection pathway which modulates their egg-laying timing. T4/T5 neurons and downstream LC10 neurons (at least partially) are involved in the neuronal circuit which detects other flies leading to female egg-laying modulation. The columnar cells T4 and T5 neurons are the first neurons to represent the direction of motion and project their axons in the different layers of the lobula plate which contain wide-field motion-sensitive neurons (Borst et al., 2020; Maisak et al., 2013; Mauss et al., 2017; Schnell et al., 2012; Zhu, 2013). Different types of LC neurons respond to either looming stimuli or motion of small objects and induce distinct behaviours such as jumping, reaching, wing extension, forward walking, backward walking, turning and result in attractive or aversive behaviours (Wu et al., 2016). More specifically, two main types of LC neurons (i.e. LC10 and LC11) encode the motion of small objects such as flies. LC10 neurons are used to track a fly-size visual object and convey visual information during courtship (Ribeiro et al., 2018). LC11 neurons are a specialized object detector and respond to movement of small objects darker than the background (Keleş and Frye, 2017). LC11 neurons are also involved in processing motion cues of other flies to downregulate freezing, movement of others being a cue of safety (Ferreira and Moita, 2020). Here we identified that LC10 neurons are involved in the detection of flies leading to a modulation of egg-laying. We however found a strong variation in egg-laying start-time between silenced-LC10 females. It can come from either an incomplete penetrance of Gal4, Gal4 not being expressed at high enough level, or by the fact that LC10 are only partially involved in group detection and other neurons participate in sensing group members in the context of egg-laying advancement.

### Interaction between light and social context

Light can impact reproduction at many levels. For instance, artificial lighting can alter timing of breeding (Navara and Nelson, 2007) and reduce egg production in birds (Asmundson et al., 1946). In German cockroaches, female’s reproductive rate is unresponsive to social interactions during the light phase, while social interactions during the dark phase are sufficient to induce faster reproduction (Uzsák and Schal, 2013), showing a potential interaction between social context and light. Our study revealed that, in *D. melanogaster*, light itself is also an important environmental parameter that interacts with the social context to modulate reproductive output in females. Light inhibits oogenesis, ovulation and egg-laying start-time only in isolated *Drosophila* females, grouped *Drosophila* females being insensitive to light conditions. Light and social environment seem therefore to be two important factors that can modify reproduction in these two species and seem to be conserved between solitary and presocial species. However, unlike German cockroaches, *Drosophila* females respond to social environment during the light phase since the presence of a group lifts the inhibition of light and stimulates egg-laying. Both species display an aversion for laying eggs under light conditions but display opposite sensitivity to social context during the light phase. This difference might be due to a stronger competition existing between *Drosophila* females than between German cockroach females, leading to a stimulation of egg-laying in the presence of others even during light phase in *Drosophila*.

Even though *Drosophila* exhibit phototactic behaviour (Zhu, 2013), *D. melanogaster* females, as other insect species (e.g. mosquitoes (Farnesi et al., 2018)), mainly lay eggs during the dark phase (Cury et al., 2019; Sheeba et al., 2001). They might act like this to avoid Ultra Violets (Zhu et al., 2014) and to protect their eggs since UV exposition leads to cellular apoptosis, damages embryos and reduction of offspring survival (Zhou and Steller, 2003). Other explanations can be that light is perceived as a stress factor by female fruit flies since the main predators of *Drosophila* (e.g. Dragonflies, parasitoid wasps) are diurnal (Carton et al., 1986; Cochard et al., 2017). The presence of light might also mean the presence of sun and a poor humidity environment leading to an increased risk of desiccation during the day. Females would therefore wait for darkness to start laying eggs to escape desiccation or other harmful effects such as parasite invasion (Howlader and Sharma, 2006), and would not in the presence of others because of competition with other females, a lower risk of desiccation (Howe, 1962b) or a dilution effect (J.L. Chapman and Reiss, 1999). Our functional experiment (Figure 1E) showed that females, during the day, escape their dish to join another dish containing a group of flies when they have the choice. We could thus predict that during the day, flies can see and spend their time trying to escape the dish in order to find a more suitable substrate next to others to lay their eggs. They might try to escape until darkness and stop at night, when they do not see anymore, which could result in a delay of egg-laying and explain why they start laying eggs at night. Females, when in a group, almost never leave the dish during the day and stay in that group for laying eggs (Figure 1E). We can suppose that females stay and lay eggs in the group to get advantages of being in a group, but do not await darkness to start laying eggs because of a strong competition between females.

### A reproductive pathway linking light and social cue detection

In our study, we explored how social environment interacts with light to modulate *D. melanogaster* female reproductive output and uncovered mechanisms that are involved in such social modulation. Based on our data, we propose the following model (Figure 7). Light inhibits and delays oogenesis in *D. melanogaster* females through the inhibition of juvenile hormone (JH) synthesis. JH level expression is then reduced in isolated females and leads to an inhibition of egg-laying. However, the presence of flies from any sex, mating status and species can lift the inhibition of light, resulting in grouped females laying eggs during the day while isolated females retain their eggs until darkness. The presence of a group is detected via visual cues through motion detection pathway and leads to egg-laying advancement that has been found to be density-dependent. The presence of others actually lifts the inhibition of light by stimulating hormonal pathway involving JH, stimulates oogenesis and leads to a faster oviposition in *Drosophila* females.

**Figure 7:**
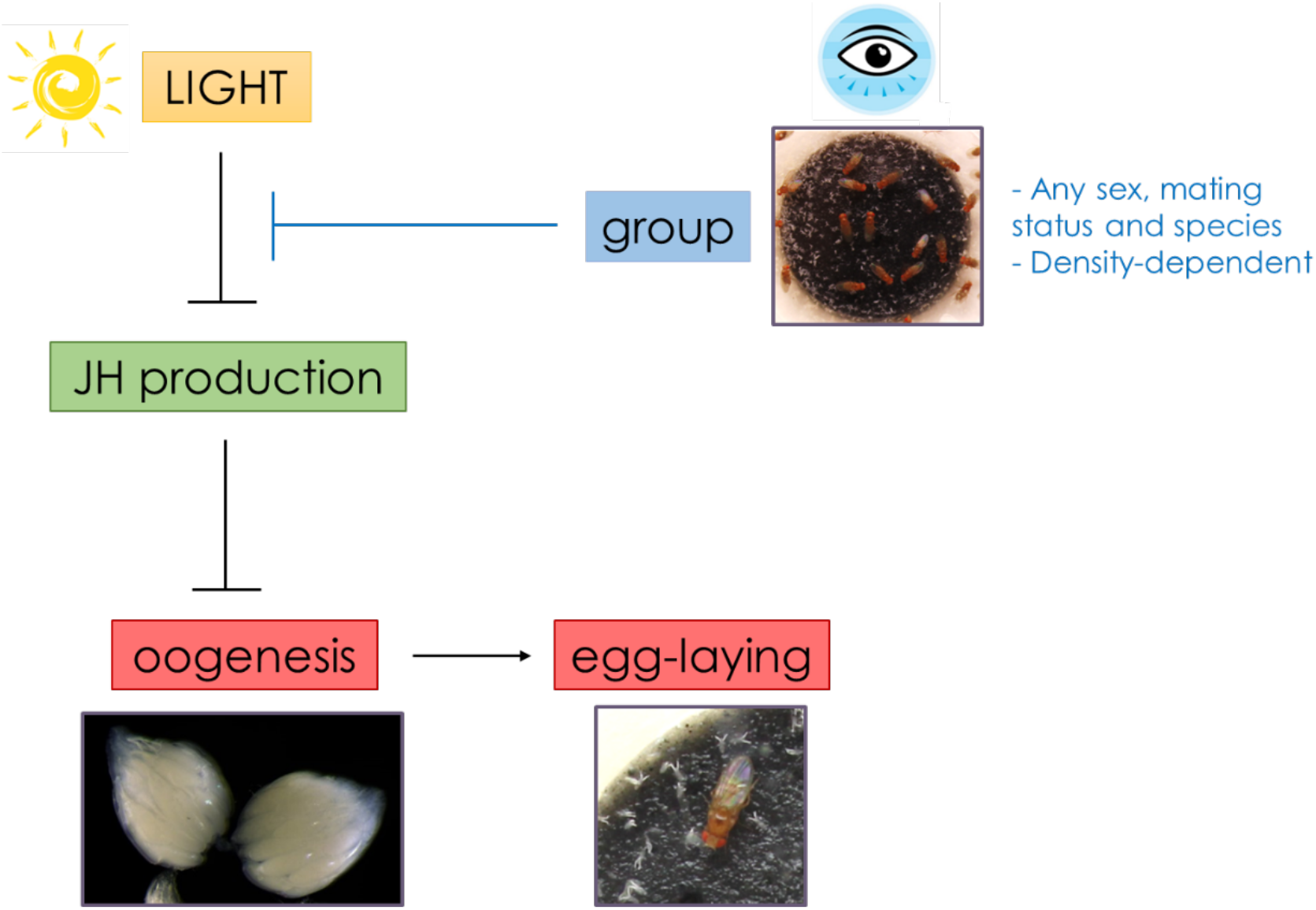
Model depicting the *D. melanogaster* female reproductive output under light condition and its modulation by the social environment.

Our findings are comparable with findings in German cockroaches, a presocial species, showing that the presence of a group modulates rate of oocyte maturation, by eliciting faster oocyte growth, and speeds up reproduction in this species (Uzsak and Schal, 2012; Uzsák and Schal, 2013). On the other side, social isolation delays oocyte development as well as sexual maturation in isolated cockroach females (Uzsak and Schal, 2012), showing that social environment modulates oogenesis the same way in this presocial species. However, despite conserved factors between these two species, input sensory modalities are different since in cockroaches, females detect the presence of others via tactile cues through antennal contacts (Uzsák and Schal, 2013), while in *Drosophila*, we showed that the presence of others is sensed by vision through motion cues. It is possible that both German cockroaches and *Drosophila* have evolved from a common ancestor for which reproduction was already modulated by social environment and then these two species diverged by using different social input sensory modality.

We further showed that application of Methoprene, a JH analog, stimulates and restores oogenesis in isolated females, leading to a similar oocyte development and oocyte rate than grouped females. This shows the involvement of juvenile hormone pathway in oocyte development. Our study suggests that light delays oogenesis in isolated females while the presence of a group lifts the inhibition of light by stimulating JH production. Social modulation of oogenesis and the involvement of JH have already been found in presocial species. In German cockroaches, social facilitation of reproduction is coupled to the endocrine system since social interactions significantly elevates the rate of JH production leading to faster oocyte development and to an acceleration of female oogenesis (Uzsak and Schal, 2012). Smaller basal oocytes correspond with lower rates of JH biosynthesis in isolated and crowded females, whereas larger and increased number of mature oocytes correspond to higher rates of JH synthesis in grouped females (Uzsak and Schal, 2012). The rate of oocyte maturation reflects the rate of JH production by the corpora allata, indicating that social interactions strongly influence the rates of JH biosynthesis in German cockroach females. Our study provides support to the ones on German cockroaches (Gadot et al., 1989; Uzsak and Schal, 2012; Uzsák and Schal, 2013) showing that groups are stimuli that affect the central nervous system which converts them into disinhibitory signals relieving JH production from brain inhibition. How visual stimuli get transduced into neuronal and neuroendocrine signals that regulate JH titer in *Drosophila* is still unknown.

Even though we found that Methoprene treatment restores oogenesis in isolated females, it did not restore ovulation and egg-laying in these females, showing that in addition of JH pathway, other neuronal and/or hormonal pathways are involved in social modulation of *Drosophila* female reproduction. For instance, another pathway might be involved at the ovulation level such as octopaminergic (OA) neurons since these neurons innervate insects’ ovary and oviducts allowing contraction of oviducts and the release of mature oocytes (Bloch Qazi et al., 2003; Li et al., 2015; Rodríguez-Valentín et al., 2006; White et al., 2021). It could explain the accumulation of mature stage 14 oocytes in ovaries of isolated females when presence of light. Further studies will need to be done to test whether the stimulation of OA neurons is able to stimulate and restore ovulation in stressed alone females. We can hypothesize that both stimulation of JH and OA neurons are necessary to restore respectively oogenesis and ovulation in isolated females and thus are both necessary to restore egg-laying in these females. These two pathways might work in combination to modulate egg-laying in female *D. melanogaster*. Our model may therefore be more complex than presented in Figure 7.

JH is synthetized by the corpora allata (CA), and stimulates insect oogenesis (Dubrovsky et al., 2002; Gujar and Palli, 2016; Hernández-Martínez et al., 2019; Santos et al., 2019; Soller et al., 1999). It has been shown that oogenesis can be affected by stressful environmental factors (e.g. food availability, presence of predators, infection with pathogenic organisms, temperature, humidity, toxicants). More precisely, stressful conditions arrest egg production in *Drosophila* via a hormonal cascade (Meiselman et al., 2018; Soller et al., 1999; Terashima et al., 2005; Terashima and Bownes, 2004) increasing ecdysone titers and indirectly resulting in reduced reproductive output through arrest of oogenesis via the inhibition of JH production (Meiselman et al., 2017, 2018; Soller et al., 1999; Terashima et al., 2005; Terashima and Bownes, 2004). Social isolation is a stress factor since social deprivation and isolation can lead to serious health problems through behavioral, psychosocial and physiological pathways in several species (Holt-Lunstad et al., 2010; Umberson and Karas Montez, 2010). For instance, social isolation causes faster mortality in ants (Koto et al., 2015), leads to very acute stress and increases size, number, distribution and malignancy of tumors in rats (Hermes et al., 2009). Isolation and reduced social interactions also lead to a faster tumor progression in *D. melanogaster* (Dawson et al., 2018). Our study reveals that JH is a modulator of reproduction and is under the control of social environment. Reduced JH level expression in isolated females might thus be due to isolation that is perceived as stressful by *Drosophila* females.

### The evolution of sociality

Females are sensitive to the presence of others in multiple species and it can affect their reproduction. As described earlier, this social modulation of reproduction entails synchronization of female ovulation in rats (McClintock, 1984; McClintock, 1981; Schank and McClintock, 1997) and a stimulation of ovulation in rabbits where females with sisters in their group display more positive social interactions, attenuating the negative consequences of stress and facilitating ovulation (Rödel et al., 2008). In eusocial insects, worker’s ovary and egg-laying are inhibited in the presence of larvae (Traynor et al., 2014; Ulrich et al., 2016) and in the presence of a queen (Hoover et al., 2003; Traynor et al., 2014). As described above, positive effect of grouping on oocyte maturation and oviposition has been found in presocial German cockroaches (Crall et al., 2016; Uzsak and Schal, 2012; Uzsák and Schal, 2013). All of these studies therefore demonstrate that the presence of a group modulates egg-laying timing in social species as well as egg-laying production, showing a social regulation of female reproductive physiology. In solitary species, social facilitation of egg-laying has been seen in *Rhagoletis pomonella* females (Prokopy and Reynolds, 1998). However, we did not find any study relating social modulation of ovarian function in solitary species. Our study therefore seems to be the first one to our knowledge showing social modulation of oogenesis, ovulation as well as egg-laying in a solitary species.

Social modulation of oogenesis and reproduction is often seen as a sign of higher sociality. Sociality is seen as an evolutionary transition from solitary to sociality, where species with complex societies differ dramatically from solitary ancestors (Kocher et al., 2014; Liu et al., 2020; Smiseth et al., 2012; Wong et al., 2013). Our study demonstrates social modulation of female reproduction in *Drosophila melanogaster*, a species that is considered solitary. Similarities in how social environment modulates female reproduction between *Drosophila melanogaster* and German cockroaches (Crall et al., 2016; Katoh et al., 2017; Uzsak and Schal, 2012; Uzsák and Schal, 2013) indicate conserved social pathways between social and solitary species, questioning the validity of the term “solitary”. We could thus suggest that sociality might have possibly evolved more as a gradient than as a transition. This further suggests that the ancestor of solitary and social species was already adjusting reproduction to social environment, meaning that, from an evolutionary perspective, social insects could have evolved from a rather competition-driven synchronization of reproduction and became highly collaborative species in part by suppressing this competition.

Our results highlight the importance of considering social environment as a main environmental factor when studying animal behaviour, even in species considered solitary. Social environment, in all species with any degree of sociality, may play an active role in modulating and creating individual variation in collective behaviours. *Drosophila* could be seen as a protosocial species containing possibilities of social evolution and could therefore be used as a model to understand the evolution of sociality.

## Materials and methods

### *Drosophila* rearing, stocks and genetics

Flies were reared on food medium (agar 10g/L, sucrose 15g/L, glucose 30g/L, dried yeast 35g/L, soy flour 10g/L, molasses 30g/L, propionic acid 5mM and tegosept 2g/L) in a 12:12h light/dark cycle at 25°C. Virgin adults were collected using CO_2_ anesthesia and aged in same-sex groups of 20 in food vials (25×95 mm) containing food medium (23×15 mm) for 5 to 7 days before testing. All focal mated females used in this study were from the wild-type *Oregon-R* strain unless stated otherwise. Different mutant lines (see Table 1) and wild-type lines (*D. simulans, D. erecta, D. suzukii, D. yakuba, D. phalerata, D. virilis*) were used as group members. Mutant and transgenic flies (see Table 1) were used as focal females to investigate sensory modalities necessary for group detection and hormonal mechanisms underlying social modulation of oogenesis and egg-laying. To remove variation arising from genetic background, all lines used as focal females were in an *Oregon-R* genetic background by 10 generations of backcrossing to an *Oregon-R* strain.

**Table 1.**
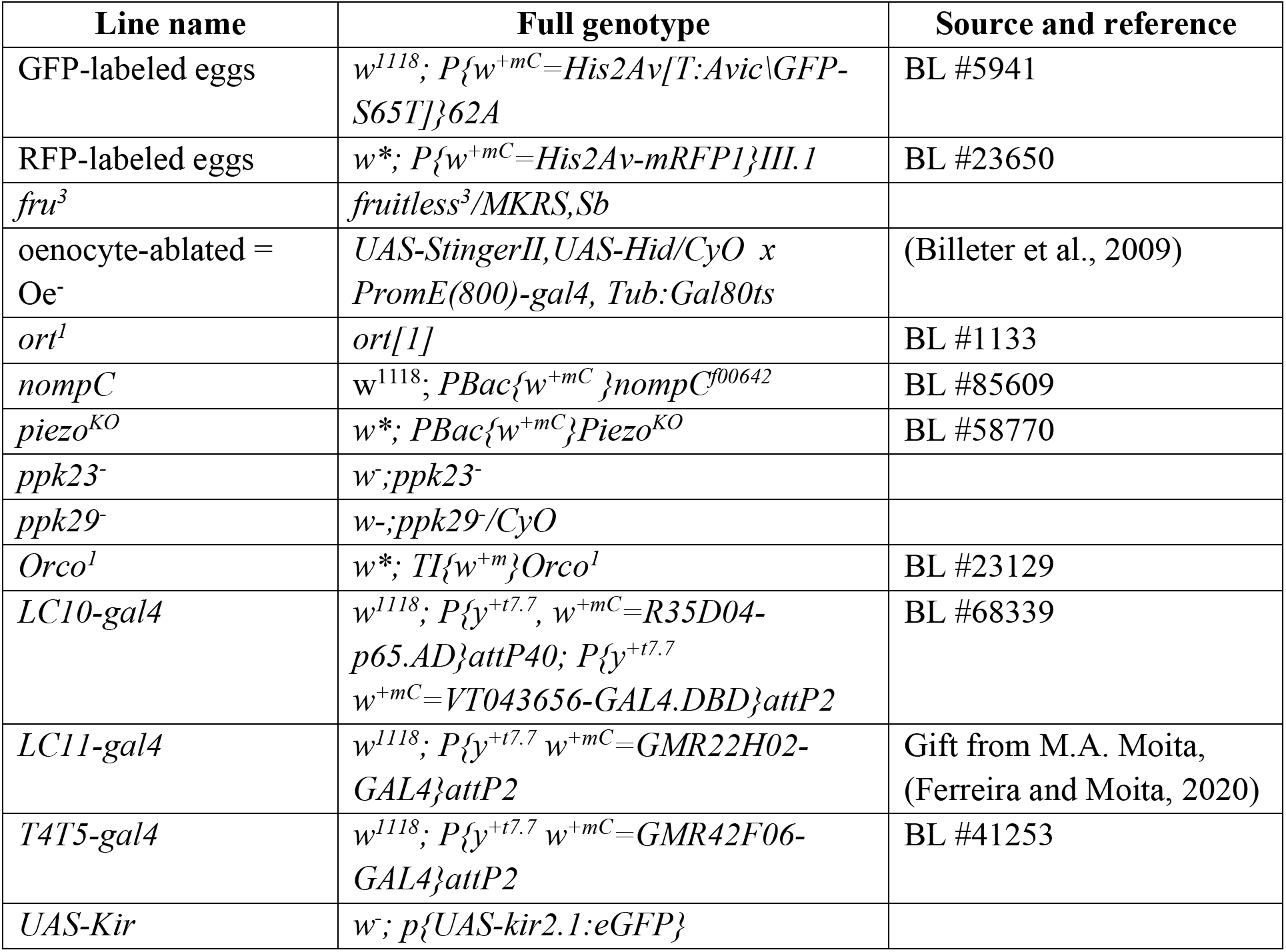
Genotypes of mutant and transgenic flies.

### Egg-laying assay

Virgin females were mated in groups of 6 females and 6 males at 25°C in mating arenas (Petri dishes of 55×14 mm containing a circular fly food patch (23×1 mm)). After 2h, females were removed and the presence of a mating plug was inspected using a UV flash light (Perel EFL41UV) to ensure females were mated (Lung and Wolfner, 2001). Mated females were placed in an egg-laying Petri dish (35×14 mm) containing a black food (food medium with 20g/L activated charcoal powder [Velda^®^]) patch (23×1mm) allowing clear visualization of the white eggs. To force females to lay eggs on the food patch surface and not on the edge, a liquid 3.5% agar solution was poured in the dish up to the level of the food patch surface. Egg-laying dishes and flies were housed in a stainless-steel enclosure [63(D) × 71(H) × 120(L) cm] for 24h at 25°C. To generate 12:12 LD condition, the chamber was lit internally by white LED lights during the light phase of the day and red (> 620 nm) LED lights during the dark phase. To quantify egg-laying, pictures of the egg-laying dishes were taken with cameras (EOS 1300D Canon, equipped with EF-S 18-55mm III lens) at 15 minutes intervals during 24h (using time-lapse software DigiCamControl). Start-time of egg-laying, number of eggs per female and cumulative number of eggs were manually determined from these pictures using software ImageJ 1.52a (Schneider et al., 2012).

### Video tracking

Bespoke fly arenas made of PETG material inspired from the FlyBowl arenas (Simon and Dickinson, 2010) were 3D printed (Creality Ender-3 V2 printer). These arenas were used to optimize video tracking of flies. Three sizes of arenas were printed (34×4mm, 59×4 mm and 94×4 mm) with the same sloped walls (gradual slope with an angle of 30° measured from the horizontal floor and with a 3mm high ceiling and 5mm length) to prevent vertical wall climbing, which can impair automated tracking. All arenas contained a circular depression in the middle (23×1 mm) in which an egg-laying food patch (23×1 mm) was placed. A thin and transparent Plexiglas plate was added on the top of arenas to avoid fly escape. After 10min habituation, a 1-hour video recording was made with raspberry Pi 4 camera module V2, allowing high quality recordings (1080p at 30fps). Groups of 6 mated females were placed in the three types of arenas and number of encounters per individual (one encounter is counted when two flies are separated by less than 3.75 mm) as well as the interindividual distance were quantified using Ethovision XT11.5 (Noldus, Wageningen, Netherlands).

### Survival assay

Females were mated as described in the egg-laying assay section. A focal wild-type mated female was placed alone or with 5 mated females laying GFP green fluorescent eggs (GFP-labelled eggs, see Table 1) in an egg-laying dish and housed as indicated in the egg-laying assay section. After 24h, females were removed from the dish and green fluorescent eggs, identified using a Leica MZ10 fluorescence stereomicroscope, were removed, keeping only the wild-type eggs laid by the focal female. Eggs from isolated and grouped *Oregon-R* females were counted in order to get the number of wild-type eggs laid in each egg-laying dish. Egg-laying dishes were maintained at 25°C in a 12:12h light/dark cycle incubator until adults enclosed, after which they were CO_2_ anesthetized and counted. Survival was calculated as the number of progenies that hatched divided by the number of wild-type eggs in the egg-laying dish.

### Food quality-dependent survival

A group of 100 mated females and 100 males was placed for 3h in a plastic square bottle (5.71cm side x 10.16cm height) containing a petri dish (35×10 mm) filled with food medium and a pinch of dried yeast in the centre for egg-laying stimulation. The petri dish was removed after 3h to collect eggs. Eggs were transferred by group (different group size according to the tested condition) on egg-laying dishes (as in egg-laying assay section). Four different food qualities were used: high quality food is composed of coal (20g/L) and the standard food (100%), medium quality food, low and very low quality food are all composed of coal (20g/L) and respectively composed of 50%, 25% and 10% of quantities of standard ingredients except for agar, propionic acid and tegosept. These dishes were maintained at 25°C in a 12:12h light/dark cycle until adults enclosed. Survival was calculated as the number of progenies that hatched divided by the number of eggs placed in the same dish at the beginning.

### Survival of host and guest eggs from homogeneous and heterogeneous groups

*Oregon-R* mated females and *W*^*1118[Oregon-R]*^ mated females (*Oregon-R* with white eyes) were used as guests and hosts, respectively. A schematic representation of the method of this assay is presented in Figure 2C. Host and guest females were mated and placed in petri dishes as described in the egg-laying assay section. The egg-laying substrate used in this assay is a medium food, composed of coal (20g/L) and 50% of quantities of standard ingredients. Host females were housed in groups of 6 and guest females were either isolated or housed in groups of 6. Fourty eggs laid by host females were kept and 10 eggs laid by guests were collected and added to the same substrate as the host eggs. Guest eggs were collected from grouped guests in synchrony with the hosts (g1) at T+4, or either from grouped guests (g2) or isolated guests (a) out of synchrony with the hosts at T+10. Homogeneous groups containing 50 eggs laid by either host or guest females were also made to control a line-related survival effect (kin effect). Petri dishes containing eggs from homogeneous and heterogeneous groups were maintained at 25°C in a 12:12h light/dark cycle until adults enclosed. Survival of host and guest eggs into viable adults was calculated for each condition as the number of progeny that hatched divided by the number of eggs placed in the same dish.

### Egg-laying and positional preference assay

Females were mated as described in the egg-laying assay section. A focal mated female was placed in an egg-laying dish connected to another egg-laying dish by a plastic tunnel (5cm length × 0.5cm diameter). The mated female can stay in the initial dish or leave it and join the other dish by crossing the tunnel. The focal mated female was either alone in the dish or with a group of 5 decapitated flies. Decapitated flies were used as group members to reduce locomotor activity and prevent crossing into the other dish. Three conditions were tested in this assay: the focal mated female started in the dish with a group of 5 decapitated flies, the second dish being socially empty (named grouped - alone); the focal mated female started in the dish alone, the second dish being also socially empty (named alone - alone); the focal mated female started in the dish alone, the second dish being with a group of 5 decapitated flies (named alone - grouped). Experimental set-ups were placed at 25°C during 24h in a 12:12h light/dark cycle. HD webcam cameras (Logitech B910 HD Webcam using the SecurityMonitor Pro software) were used to continuously record the location of the focal mated female. Percentage of leaving and the latency to leave the initial dish as well as the time spent in each dish, were quantified by visual inspection of the videos.

### Pheromonal extraction

Pheromones were extracted and applied as in Dumenil et al., 2016 (Duménil et al., 2016b). 5 to 7 days after eclosion, males and mated females were frozen at −20°C for minimum 1 hour and separated in sex specific groups. For each group, the number of flies was counted and placed in 25mL vial with hexane. Due to evaporation and absorption of hexane, an extra 5% hexane volume is added, giving a volume of 12μL hexane per fly. After vortexing 3min, the supernatant contained pheromonal extracts of flies and were used for experiments. Whatman filter paper discs (Whatman CHR1) of 6 mm diameter together with parafilm were cut and 5 of them were placed around the food patch. 10μL of pheromonal extract were added on each filter paper to mimic as much as possible the smell of the presence of 5 group members.

### Mirror assay

A flexible and adhesive mirror (Bangood, ID: 1355324) was cut in small pieces (11cm length × 1cm width) and applied along the inside part of the egg-laying assay dishes (35×14 mm) in order to cover all the circular inside dish walls. Egg-laying assays were performed in petri dishes containing the circular mirror as described in the egg-laying assay section.

### Anti-UVs filter

Anti-UVs filters were used to block UVs below 400nm (Rosco laboratories, product number: 101031142024). Three sheets of filter were placed side by side below the LED lights inside the experimental chamber where egg-laying assays are performed as described in the egg-laying assay section.

### Ovaries dissection

Females were mated as described in the egg-laying assay section. Ovaries from isolated and grouped mated females that were housed in a 12:12 light:dark cycle or in a full dark cycle were dissected 2h and 6h after mating. To do so, females were anesthetized with CO_2_ and then sacrificed by immersion in a 70% ethanol bath. Ovaries and oviduct were dissected in 1x PBS with tweezers (Dumont #5, Fine Science Tools Ltd). Percentage of females that are ovulating was determined by the presence or absence of an egg in the lateral oviducts. After isolation, ovaries were fixed in 4% formaldehyde during 30 minutes and then washed three times in 1x PBS. Ovarioles were dissected to release oocytes and to be able to count them. Number of oocytes per ovary for oocyte stages 9 to 14 (14 being the more mature oocyte stage) were counted under a stereomicroscope (Leica MZ10F). To quantify the total number of mature eggs per female, the number of eggs laid after mating and before dissection, the number of eggs in oviduct and the number of eggs at stage 14 in ovaries were counted. The experimenter was blinded to the genotype and treatment.

### Methoprene treatment

0.5% Methoprene solution was prepared by diluting 1μL Methoprene (Methoprene Pestanal^®^ Sigma-Aldrich) in 200μL of 100% Acetone. Flies were mated as described in the egg-laying assay section. After mating, females were anaesthetized on ice and treated with Methoprene by directly applying 1μL of solution on the abdomen of the females by using a micropipette (Wijesekera et al., 2016). Control flies were treated with 1μL of pure Acetone. After 15 minutes recovery, females treated with Methoprene or Acetone were placed individually in egg-laying dishes. Pictures of egg-laying dishes were taken and analysed as described in the egg-laying assay section.

### Statistical analysis

Statistical analyses were performed using R 3.5.0 software (The R Foundation for Statistical Computing). Residuals were visually inspected for normality, and homogeneity of variances were evaluated with the Levene’s test. The Wilcoxon-Mann-Whitney test and Student’s t-test were used to compare two groups (alone vs grouped conditions). The Kruskal-Wallis test followed by post-hoc comparisons (with Bonferroni correction) and the Anova test followed by Tukey post-hoc comparisons were used to compare more than two groups. A zero-inflated linear model (fitted with a binomial distribution) was used to analyse multiple comparisons in egg survival according to group conditions. Generalized linear models (with binomial distribution) were used to analyse multiple comparisons in dish preference and index preference according to social conditions. A generalized linear model (with Poisson distribution corrected for overdispersion) was used to analyse multiple comparisons in tunnel crosses according to social conditions. Interactions and main effects of factors were determined by a type 3 two-way ANOVA, followed by post-hoc comparisons of the interaction (with Holm adjustment method). All models were then followed by Tukey post-hoc comparisons.

All graphs and some statistical analyses were performed using GraphPad Prism 9.0.0 software. Linear regressions were calculated to see correlations between egg-laying start-time and density, between number of contacts and density and between inter-individual distance and density. Non-linear regression (exponential type) was also calculated to see correlation between egg-laying start-time and density. The two-way repeated-measurements Anova, followed by post-hoc comparisons (with Sidak’s correction) were used to analyze dynamics of egg-laying between alone and grouped conditions.

## Acknowledgements

We thank Michael van Dijk for designing 3D Flybowls and Sanne Lamers for help with the video tracking. We thank Audrey Bailly and Adithya Sarma for their help during the experiments in Fig. 3A and Fig. 3C, as well as Tom Sarraude for his help with the statistics. We thank Carla Ferreira and Marta Moita for generously sharing fly stocks and Ines Ribeiro and Marion Silies for help with the visual pathway experiments. This work was funded by an adaptive life scholarship from the Univeristy of Groningen to JCB, BW, RE and TPMB.

## Notes

### Competing Interest Statement

The authors have declared no competing interest.

## References

Agrawal, S., Safarik, S., Dickinson, M.H., 2014. The relative roles of vision and chemosensation in mate recognition of *Drosophila*. Journal of Experimental Biology jeb.105817. https://doi.org/10.1242/jeb.105817

Ali, A., 2016. Effect of UV-A radiation as an environmental stress on the development, longevity, and reproduction of the oriental armyworm, Mythimna separata (Lepidoptera: Noctuidae). Environ Sci Pollut Res 6.

Allee, W.C., 1927a. Animal Aggregations. The Quarterly Review of Biology 2, 367–398.

Allee, W.C., 1927b. Animal Aggregations. The Quarterly Review of Biology 2, 367–398.

Amrein, H., 2004. Pheromone perception and behavior in. Current Opinion in Neurobiology 14, 435–442. https://doi.org/10.1016/j.conb.2004.07.008

Bartelt, R.J., Schaner, A.M., Jackson, L.L., 1985. cis-Vaccenyl acetate as an aggregation pheromone inDrosophila melanogaster. J Chem Ecol 11, 1747–1756. https://doi.org/10.1007/BF01012124

Bidaye, S.S., Laturney, M., Chang, A.K., Liu, Y., Bockemühl, T., Büschges, A., Scott, K., 2020. Two Brain Pathways Initiate Distinct Forward Walking Programs in Drosophila. Neuron 108, 469–485.e8. https://doi.org/10.1016/j.neuron.2020.07.032

Billeter, J.-C., Atallah, J., Krupp, J.J., Millar, J.G., Levine, J.D., 2009. Specialized cells tag sexual and species identity in Drosophila melanogaster. Nature 461, 987–991. https://doi.org/10.1038/nature08495

Billeter, J.-C., Jagadeesh, S., Stepek, N., Azanchi, R., Levine, J.D., 2012. Drosophila melanogaster females change mating behaviour and offspring production based on social context. Proceedings of the Royal Society B: Biological Sciences 279, 2417–2425. https://doi.org/10.1098/rspb.2011.2676

Billeter, J.-C., Levine, J.D., 2015. The role of cVA and the Odorant binding protein Lush in social and sexual behavior in Drosophila melanogaster. Frontiers in Ecology and Evolution 3. https://doi.org/10.3389/fevo.2015.00075

Billeter, J.-C., Levine, J.D., 2013. Who is he and what is he to you? Recognition in Drosophila melanogaster. Current Opinion in Neurobiology 23, 17–23. https://doi.org/10.1016/j.conb.2012.08.009

Billeter, J.-C., Wolfner, M.F., 2018. Chemical Cues that Guide Female Reproduction in Drosophila melanogaster. J Chem Ecol 44, 750–769. https://doi.org/10.1007/s10886-018-0947-z

Bloch Qazi, M.C., Heifetz, Y., Wolfner, M.F., 2003. The developments between gametogenesis and fertilization: ovulation and female sperm storage in drosophila melanogaster. Developmental Biology 256, 195–211. https://doi.org/10.1016/S0012-1606(02)00125-2

Borst, A., Haag, J., Mauss, A.S., 2020. How fly neurons compute the direction of visual motion. J Comp Physiol A 206, 109–124. https://doi.org/10.1007/s00359-019-01375-9

Brown, J.L., Sheffield, D., Leary, M.R., Robinson, M.E., 2003. Social Support and Experimental Pain. Psychosomatic Medicine 65, 276. https://doi.org/10.1097/01.PSY.0000030388.62434.46

Carton, Y., Boulétreau, M., Van Alphen, J.J.M., Van Lenteren, J.C., 1986. The Drosophila Parasitic Wasps. London, UK: Academic Press The genetics and biology of Drosophila, 347–394.

Chapman, J.L., Reiss, M., 1999. Ecology : Principles and Applications, 2nd edition Cambridge: Cambridge University Press. ed.

Chapman, J. L., Reiss, M.J., 1999. Ecology: Principles and Applications. Cambridge University Press.

Chyb, S., Gompel, N., 2013. D. melanogaster subgroup species, in: Atlas of Drosophila Morphology. Elsevier, pp. 209–220. https://doi.org/10.1016/B978-0-12-384688-4.00010-9

Clutton-Brock, T., Huchard, E., 2013. Social competition and its consequences in female mammals: Female reproductive competition in mammals. J Zool 289, 151–171. https://doi.org/10.1111/jzo.12023

Clutton-Brock, T.H., 1991. The evolution of parental care, Monographs in behavior and ecology. Princeton University Press, Princeton, N.J.

Cochard, P., Galstian, T., Cloutier, C., 2017. Light Environments Differently Affect Parasitoid Wasps and their Hosts’ Locomotor Activity. J Insect Behav 30, 595–611. https://doi.org/10.1007/s10905-017-9644-y

Coste, B., Xiao, B., Santos, J.S., Syeda, R., Grandl, J., Spencer, K.S., Kim, S.E., Schmidt, M., Mathur, J., Dubin, A.E., Montal, M., Patapoutian, A., 2012. Piezo proteins are pore-forming subunits of mechanically activated channels. Nature 483, 176–181. https://doi.org/10.1038/nature10812

Courchamp, F., Clutton-Brock, T., Grenfell, B., 1999. Inverse density dependence and the Allee effect. Trends in Ecology & Evolution 14, 405–410. https://doi.org/10.1016/S0169-5347(99)01683-3

Crall, J.D., Souffrant, A.D., Akandwanaho, D., Hescock, S.D., Callan, S.E., Coronado, W.M., Baldwin, M.W., de Bivort, B.L., 2016. Social context modulates idiosyncrasy of behaviour in the gregarious cockroach Blaberus discoidalis. Animal Behaviour 111, 297–305. https://doi.org/10.1016/j.anbehav.2015.10.032

Cury, K.M., Prud’homme, B., Gompel, N., 2019. A short guide to insect oviposition: when, where and how to lay an egg. Journal of Neurogenetics 33, 75–89. https://doi.org/10.1080/01677063.2019.1586898

Davies, L., 1998. **Delayed egg production and a possible group effect in the blowfly *Calliphora vicina*** : Delayed egg production in C. vicina. Medical and Veterinary Entomology 12, 339–344. https://doi.org/10.1046/j.1365-2915.1998.00124.x

Dawson, E.H., Bailly, T.P.M., Dos Santos, J., Moreno, C., Devilliers, M., Maroni, B., Sueur, C., Casali, A., Ujvari, B., Thomas, F., Montagne, J., Mery, F., 2018. Social environment mediates cancer progression in Drosophila. Nature Communications 9. https://doi.org/10.1038/s41467-018-05737-w

DeLong, K.T., 1978. The Effect of the Manipulation of Social Structure on Reproduction in House Mice. Ecology 59, 922–933. https://doi.org/10.2307/1938544

Demir, E., Dickson, B.J., 2005. fruitless Splicing Specifies Male Courtship Behavior in Drosophila. Cell 121, 785–794. https://doi.org/10.1016/j.cell.2005.04.027

Drickamer, L.C., 1983. Male Acceleration of Puberty in Female Mice (Mus musculus). Journal of Comparative Psychology 97, 191–200.

Dubrovsky, E.B., Dubrovskaya, V.A., Berger, E.M., 2002. Juvenile hormone signaling during oogenesis in Drosophila melanogaster. Insect Biochemistry and Molecular Biology 32, 1555–1565. https://doi.org/10.1016/S0965-1748(02)00076-0

Duménil, C., Woud, D., Pinto, F., Alkema, J.T., Jansen, I., Van Der Geest, A.M., Roessingh, S., Billeter, J.-C., 2016a. Pheromonal Cues Deposited by Mated Females Convey Social Information about Egg-Laying Sites in Drosophila Melanogaster. Journal of Chemical Ecology 42, 259–269. https://doi.org/10.1007/s10886-016-0681-3

Duménil, C., Woud, D., Pinto, F., Alkema, J.T., Jansen, I., Van Der Geest, A.M., Roessingh, S., Billeter, J.-C., 2016b. Pheromonal Cues Deposited by Mated Females Convey Social Information about Egg-Laying Sites in Drosophila Melanogaster. J Chem Ecol 42, 259–269. https://doi.org/10.1007/s10886-016-0681-3

Dunbar R. I. M, Dunbar E. P., 1977. Dominance and reproductive success among female gelada baboons. Nature 351–352.

Etienne, R., Wertheim, B., Hemerik, L., Schneider, P., Powell, J., 2002a. The interaction between dispersal, the Allee effect and scramble competition affects population dynamics. Ecological Modelling 148, 153–168. https://doi.org/10.1016/S0304-3800(01)00417-3

Etienne, R., Wertheim, B., Hemerik, L., Schneider, P., Powell, J., 2002b. The interaction between dispersal, the Allee effect and scramble competition affects population dynamics. Ecological Modelling 148, 153–168. https://doi.org/10.1016/S0304-3800(01)00417-3

Everaerts, C., Farine, J.-P., Cobb, M., Ferveur, J.-F., 2010. Drosophila Cuticular Hydrocarbons Revisited: Mating Status Alters Cuticular Profiles. PLoS ONE 5, e9607. https://doi.org/10.1371/journal.pone.0009607

Farnesi, L.C., Barbosa, C.S., Araripe, L.O., Bruno, R.V., 2018. The influence of a light and dark cycle on the egg laying activity of Aedes aegypti (Linnaeus, 1762) (Diptera: Culicidae). Mem. Inst. Oswaldo Cruz 113. https://doi.org/10.1590/0074-02760170362

Faruki, S.I., Das, D.R., Khan, A.R., Khatun, M., 2007. Effects of Ultraviolet (254nm) Irradiation on Egg Hatching and Adult Emergence of the Flour Beetles, *Tribolium castaneum*, *T. confusum* and the Almond Moth, *Cadra cautella*. Journal of Insect Science 7, 1–6. https://doi.org/10.1673/031.007.3601

Ferreira, C.H., Moita, M.A., 2020. Behavioral and neuronal underpinnings of safety in numbers in fruit flies. Nat Commun 11, 4182. https://doi.org/10.1038/s41467-020-17856-4

Ferveur, J.-F., 2005. Cuticular Hydrocarbons: Their Evolution and Roles in Drosophila Pheromonal Communication. Behav Genet 35, 279–295. https://doi.org/10.1007/s10519-005-3220-5

Ferveur, J.-F., 1997. The pheromonal role of cuticular hydrocarbons inDrosophila melanogaster. Bioessays 19, 353–358. https://doi.org/10.1002/bies.950190413

Frank, S.A., 2003. REPRESSION OF COMPETITION AND THE EVOLUTION OF COOPERATION. Evolution 57, 693–705. https://doi.org/10.1111/j.0014-3820.2003.tb00283.x

Gadot, M., Burns, E., Schal, C., 1989. Juvenile hormone biosynthesis and oocyte development in adult femaleBlattella germanica: Effects of grouping and mating. Arch. Insect Biochem. Physiol. 11, 189–200. https://doi.org/10.1002/arch.940110306

Gengs, C., Leung, H.-T., Skingsley, D.R., Iovchev, M.I., Yin, Z., Semenov, E.P., Burg, M.G., Hardie, R.C., Pak, W.L., 2002. The Target of Drosophila Photoreceptor Synaptic Transmission Is a Histamine-gated Chloride Channel Encoded byort (hclA). Journal of Biological Chemistry 277, 42113–42120. https://doi.org/10.1074/jbc.M207133200

Gorter, J.A., Jagadeesh, S., Gahr, C., Boonekamp, J.J., Levine, J.D., Billeter, J.-C., 2016. The nutritional and hedonic value of food modulate sexual receptivity in Drosophila melanogaster females. Scientific Reports 6. https://doi.org/10.1038/srep19441

Gujar, H., Palli, S.R., 2016. Juvenile hormone regulation of female reproduction in the common bed bug, Cimex lectularius. Sci Rep 6, 35546. https://doi.org/10.1038/srep35546

Hall, J.C., 1994. The Mating of a Fly. Science, New Series 264, 1702–1714.

Hermes, G.L., Delgado, B., Tretiakova, M., Cavigelli, S.A., Krausz, T., Conzen, S.D., McClintock, M.K., 2009. Social isolation dysregulates endocrine and behavioral stress while increasing malignant burden of spontaneous mammary tumors. PNAS 106, 22393–22398. https://doi.org/10.1073/pnas.0910753106

Hernández-Martínez, S., Cardoso-Jaime, V., Nouzova, M., Michalkova, V., Ramirez, C.E., Fernandez-Lima, F., Noriega, F.G., 2019. Juvenile hormone controls ovarian development in female Anopheles albimanus mosquitoes. Sci Rep 9, 2127. https://doi.org/10.1038/s41598-019-38631-6

Holt-Lunstad, J., Smith, T.B., Layton, J.B., 2010. Social Relationships and Mortality Risk: A Meta-Analytic Review.

Hoover, S.E.R., Keeling, C.I., Winston, M.L., Slessor, K.N., 2003. The effect of queen pheromones on worker honey bee ovary development. Naturwissenschaften 90, 477–480. https://doi.org/10.1007/s00114-003-0462-z

Howe, R.W., 1962a. A study of the heating of stored grain caused by insects. Annals of Applied Biology 50, 137–158. https://doi.org/10.1111/j.1744-7348.1962.tb05995.x

Howe, R.W., 1962b. A study of the heating of stored grain caused by insects. Ann Applied Biology 50, 137–158. https://doi.org/10.1111/j.1744-7348.1962.tb05995.x

Howlader, G., Sharma, V.K., 2006. Circadian regulation of egg-laying behavior in fruit flies Drosophila melanogaster. Journal of Insect Physiology 52, 779–785. https://doi.org/10.1016/j.jinsphys.2006.05.001

Jallon, J.-M., 1984. A few chemical words exchanged byDrosophila during courtship and mating. Behav Genet 14, 441–478. https://doi.org/10.1007/BF01065444

Kacsoh, B.Z., Bozler, J., Ramaswami, M., Bosco, G., 2015. Social communication of predator-induced changes in Drosophila behavior and germ line physiology. eLife 4, e07423. https://doi.org/10.7554/eLife.07423

Katoh, K., Iwasaki, M., Hosono, S., Yoritsune, A., Ochiai, M., Mizunami, M., Nishino, H., 2017. Group-housed females promote production of asexual ootheca in American cockroaches. Zoological Lett 3, 3. https://doi.org/10.1186/s40851-017-0063-x

Keleş, M.F., Frye, M.A., 2017. Object-Detecting Neurons in Drosophila. Current Biology 27, 680–687. https://doi.org/10.1016/j.cub.2017.01.012

Kim, W.J., Jan, L.Y., Jan, Y.N., 2012. Contribution of visual and circadian neural circuits to memory for prolonged mating induced by rivals. Nat Neurosci 15, 876–883. https://doi.org/10.1038/nn.3104

Kocher, S.D., Pellissier, L., Veller, C., Purcell, J., Nowak, M.A., Chapuisat, M., Pierce, N.E., 2014. Transitions in social complexity along elevational gradients reveal a combined impact of season length and development time on social evolution. Proc. R. Soc. B. 281, 20140627. https://doi.org/10.1098/rspb.2014.0627

Koto, A., Mersch, D., Hollis, B., Keller, L., 2015. Social isolation causes mortality by disrupting energy homeostasis in ants. Behav Ecol Sociobiol 69, 583–591. https://doi.org/10.1007/s00265-014-1869-6

Krupp, J.J., Kent, C., Billeter, J.-C., Azanchi, R., So, A.K.-C., Schonfeld, J.A., Smith, B.P., Lucas, C., Levine, J.D., 2008. Social Experience Modifies Pheromone Expression and Mating Behavior in Male Drosophila melanogaster. Current Biology 18, 1373–1383. https://doi.org/10.1016/j.cub.2008.07.089

Larsson, M.C., Domingos, A.I., Jones, W.D., Chiappe, M.E., Amrein, H., Vosshall, L.B., 2004. Or83b Encodes a Broadly Expressed Odorant Receptor Essential for Drosophila Olfaction. Neuron 43, 703–714. https://doi.org/10.1016/j.neuron.2004.08.019

Laturney, M., Billeter, J.-C., 2016. Drosophila melanogaster females restore their attractiveness after mating by removing male anti-aphrodisiac pheromones. Nat Commun 7, 12322. https://doi.org/10.1038/ncomms12322

Laws, R.M., 1929. Aspects of reproduction in the African elephant, Loxodonta africana. J. Reprod. Fertil. Suppl. 6, 193–218.

Le Conte, Y., Mohammedi, A., Robinson, G.E., 2001. Primer effects of a brood pheromone on honeybee behavioural development. Proc. R. Soc. Lond. B 268, 163–168. https://doi.org/10.1098/rspb.2000.1345

Li, Y., Fink, C., El-Kholy, S., Roeder, T., 2015. THE OCTOPAMINE RECEPTOR octß2R IS ESSENTIAL FOR OVULATION AND FERTILIZATION IN THE FRUIT FLY *Drosophila melanogaster*: octß2R is essential for ovulation. Arch. Insect Biochem. Physiol. 88, 168–178. https://doi.org/10.1002/arch.21211

Lihoreau, M., Brepson, L., Rivault, C., 2009. The weight of the clan: Even in insects, social isolation can induce a behavioural syndrome. Behavioural Processes 82, 81–84. https://doi.org/10.1016/j.beproc.2009.03.008

Liu (劉彥廷), M., Chan (詹仕凡), S.-F., Rubenstein, D.R., Sun (孫烜駿), S.-J., Chen (陳伯飛), B.-F., Shen (沈聖峰), S.-F., 2020. Ecological Transitions in Grouping Benefits Explain the Paradox of Environmental Quality and Sociality. The American Naturalist 195, 818–832. https://doi.org/10.1086/708185

Lu, B., LaMora, A., Sun, Y., Welsh, M.J., Ben-Shahar, Y., 2012. ppk23-Dependent Chemosensory Functions Contribute to Courtship Behavior in Drosophila melanogaster. PLoS Genet 8, e1002587. https://doi.org/10.1371/journal.pgen.1002587

Lung, O., Wolfner, M.F., 2001. Identification and characterization of the major Drosophila melanogaster mating plug protein. Insect Biochemistry and Molecular Biology 9.

Maisak, M.S., Haag, J., Ammer, G., Serbe, E., Meier, M., Leonhardt, A., Schilling, T., Bahl, A., Rubin, G.M., Nern, A., Dickson, B.J., Reiff, D.F., Hopp, E., Borst, A., 2013. A directional tuning map of Drosophila elementary motion detectors. Nature 500, 212–216. https://doi.org/10.1038/nature12320

Manjunatha, T., Dass, S.H., Sharma, V.K., 2008. Egg-laying rhythm in Drosophila melanogaster. J Genet 87, 495–504. https://doi.org/10.1007/s12041-008-0072-9

Mauss, A.S., Vlasits, A., Borst, A., Feller, M., 2017. Visual Circuits for Direction Selectivity. Annu. Rev. Neurosci. 40, 211–230. https://doi.org/10.1146/annurev-neuro-072116-031335

McClintock, M., 1984. Estrous synchrony: Modulation of ovarian cycle length by female pheromones. Physiology & Behavior 32, 701–705. https://doi.org/10.1016/0031-9384(84)90181-1

Mcclintock, M.K., 1981. Social Control of the Ovarian Cycle and the Function of Estrous Synchrony. Am Zool 21, 243–256. https://doi.org/10.1093/icb/21.1.243

Meiselman, M., Lee, S.S., Tran, R.-T., Dai, H., Ding, Y., Rivera-Perez, C., Wijesekera, T.P., Dauwalder, B., Noriega, F.G., Adams, M.E., 2017. Endocrine network essential for reproductive success in *Drosophila melanogaster*. Proc Natl Acad Sci USA 114, E3849–E3858. https://doi.org/10.1073/pnas.1620760114

Meiselman, M.R., Kingan, T.G., Adams, M.E., 2018. Stress-induced reproductive arrest in Drosophila occurs through ETH deficiency-mediated suppression of oogenesis and ovulation. BMC Biol 16, 18. https://doi.org/10.1186/s12915-018-0484-9

Meng, J.-Y., Zhang, C.-Y., Zhu, F., Wang, X.-P., Lei, C.-L., 2009. Ultraviolet light-induced oxidative stress: Effects on antioxidant response of Helicoverpa armigera adults. Journal of Insect Physiology 55, 588–592. https://doi.org/10.1016/j.jinsphys.2009.03.003

Mohammedi, A., Paris, A., Crauser, D., Le Conte, Y., 1998. Effect of Aliphatic Esters on Ovary Development of Queenless Bees ( Apis mellifera L.). Naturwissenschaften 85, 455–458. https://doi.org/10.1007/s001140050531

Muller-Schwarze, D., 1984. *Pheromones and Reproduction in Mammals.* John G. Vandenbergh. The Quarterly Review of Biology 59, 325–325. https://doi.org/10.1086/413940

Narasimha, S., Nagornov, K.O., Menin, L., Mucciolo, A., Rohwedder, A., Humbel, B.M., Stevens, M., Thum, A.S., Tsybin, Y.O., Vijendravarma, R.K., 2019. Drosophila melanogaster cloak their eggs with pheromones, which prevents cannibalism. PLoS Biol 17, e2006012. https://doi.org/10.1371/journal.pbio.2006012

Navara, K.J., Nelson, R.J., 2007. The dark side of light at night: physiological, epidemiological, and ecological consequences. J Pineal Res 43, 215–224. https://doi.org/10.1111/j.1600-079X.2007.00473.x

Oldroyd, B., Wossler, T., Ratnieks, F., 2001. Regulation of ovary activation in worker honey-bees ( Apis mellifera ): larval signal production and adult response thresholds differ between anarchistic and wild-type bees. Behavioral Ecology and Sociobiology 50, 366–370. https://doi.org/10.1007/s002650100369

Peyser MR, Ayalon D, Harell A, Toaff R, Cardova T, 1973. Stress induced delay of ovulation. Obstet Gynecol. 667–71. PMID: 4749568.

Prokopy, R.J., Reynolds, A.H., 1998. Ovipositional enhancement through socially facilitated behavior in Rhagoletis pomonella flies. Entomologia Experimentalis et Applicata 86, 281–286. https://doi.org/10.1046/j.1570-7458.1998.00290.x

Ribeiro, I.M.A., Drews, M., Bahl, A., Machacek, C., Borst, A., Dickson, B.J., 2018. Visual Projection Neurons Mediating Directed Courtship in Drosophila. Cell 174, 607–621.e18. https://doi.org/10.1016/j.cell.2018.06.020

Richard, D.S., Watkins, N.L., Serafin, R.B., Gilbert, L.I., 1998. Ecdysteroids regulate yolk protein uptake by Drosophila melanogaster oocytes. Journal of Insect Physiology 44, 637–644. https://doi.org/10.1016/S0022-1910(98)00020-1

Rödel, H.G., Starkloff, A., Bruchner, B., von Holst, D., 2008. Social environment and reproduction in female European rabbits (Oryctolagus cuniculus): Benefits of the presence of litter sisters. Journal of Comparative Psychology 122, 73–83. https://doi.org/10.1037/0735-7036.122.1.73

Rodríguez-Valentín, R., López-González, I., Jorquera, R., Labarca, P., Zurita, M., Reynaud, E., 2006. Oviduct contraction in *Drosophila* is modulated by a neural network that is both, octopaminergic and glutamatergic. J. Cell. Physiol. 209, 183–198. https://doi.org/10.1002/jcp.20722

Rooke, R., Rasool, A., Schneider, J., Levine, J.D., 2020. Drosophila melanogaster behaviour changes in different social environments based on group size and density. Commun Biol 3, 304. https://doi.org/10.1038/s42003-020-1024-z

Rowell, T.E., 1970. Baboon menstrual cycles affected by social environment. J Reprod Fertil. (1):133–41. doi: 10.1530/jrf.0.0210133. PMID: 4983966.

Sakata, H., Katayama, N., 2001. Ant defence system: A mechanism organizing individual responses into efficient collective behavior: Ant defence system. Ecological Research 16, 395–403. https://doi.org/10.1046/j.1440-1703.2001.00404.x

Santos, C.G., Humann, F.C., Hartfelder, K., 2019. Juvenile hormone signaling in insect oogenesis. Current Opinion in Insect Science 31, 43–48. https://doi.org/10.1016/j.cois.2018.07.010

Schank, J.C., McClintock, M.K., 1997. Ovulatory Pheromone Shortens Ovarian Cycles of Female Rats Living in Olfactory Isolation 6.

Schneider, C.A., Rasband, W.S., Eliceiri, K.W., 2012. NIH Image to ImageJ: 25 years of image analysis. Nat Methods 9, 671–675. https://doi.org/10.1038/nmeth.2089

Schnell, B., Raghu, S.V., Nern, A., Borst, A., 2012. Columnar cells necessary for motion responses of wide-field visual interneurons in Drosophila. J Comp Physiol A 198, 389–395. https://doi.org/10.1007/s00359-012-0716-3

Scott, D., 1994. GENETIC VARIATION FOR FEMALE MATE DISCRIMINATION IN *DROSOPHILA MELANOGASTER*. Evolution 48, 112–121. https://doi.org/10.1111/j.1558-5646.1994.tb01298.x

Seeholzer, L.F., Seppo, M., Stern, D.L., Ruta, V., 2018. Evolution of a central neural circuit underlies Drosophila mate preferences. Nature 559, 564–569. https://doi.org/10.1038/s41586-018-0322-9

Sevenster, J.G., 1996. Aggregation and Coexistence. I. Theory and Analysis. The Journal of Animal Ecology 65, 297. https://doi.org/10.2307/5876

Sheeba, V., Chandrashekaran, M.K., Joshi, A., Kumar Sharma, V., 2001. A case for multiple oscillators controlling different circadian rhythms in Drosophila melanogaster. Journal of Insect Physiology 47, 1217–1225. https://doi.org/10.1016/S0022-1910(01)00107-X

Silbering, A.F., Rytz, R., Grosjean, Y., Abuin, L., Ramdya, P., Jefferis, G.S.X.E., Benton, R., 2011. Complementary Function and Integrated Wiring of the Evolutionarily Distinct Drosophila Olfactory Subsystems. Journal of Neuroscience 31, 13357–13375. https://doi.org/10.1523/JNEUROSCI.2360-11.2011

Simon, J.C., Dickinson, M.H., 2010. A New Chamber for Studying the Behavior of Drosophila. PLoS ONE 5, e8793. https://doi.org/10.1371/journal.pone.0008793

Smiseth, P.T., Kölliker, M., Royle, N.J., 2012. What is parental care?, in: Royle, N.J., Smiseth, P.T. (Eds.), The Evolution of Parental Care. Oxford University Press, pp. 1–17. https://doi.org/10.1093/acprof:oso/9780199692576.003.0001

Soller, M., Bownes, M., Kubli, E., 1999. Control of Oocyte Maturation in Sexually MatureDrosophilaFemales. Developmental Biology 208, 337–351. https://doi.org/10.1006/dbio.1999.9210

Stephens, P.A., Sutherland, W.J., 1999. Consequences of the Allee effect for behaviour, ecology and conservation. Trends in Ecology & Evolution 14, 401–405. https://doi.org/10.1016/S0169-5347(99)01684-5

Stockley, P., Bro-Jørgensen, J., 2011. Female competition and its evolutionary consequences in mammals. Biological Reviews 86, 341–366. https://doi.org/10.1111/j.1469-185X.2010.00149.x

Symonds, M.R.E., Wertheim, B., 2005. The mode of evolution of aggregation pheromones in Drosophila species: Pheromone evolution in Drosophila. Journal of Evolutionary Biology 18, 1253–1263. https://doi.org/10.1111/j.1420-9101.2005.00971.x

Taborsky, B., Skubic, E., Bruintjes, R., 2007. Mothers adjust egg size to helper number in a cooperatively breeding cichlid. Behavioral Ecology 18, 652–657. https://doi.org/10.1093/beheco/arm026

Terashima, J., Bownes, M., 2004. Translating Available Food Into the Number of Eggs Laid by *Drosophila melanogaster*. Genetics 167, 1711–1719. https://doi.org/10.1534/genetics.103.024323

Terashima, J., Takaki, K., Sakurai, S., Bownes, M., 2005. Nutritional status affects 20-hydroxyecdysone concentration and progression of oogenesis in Drosophila melanogaster. Journal of Endocrinology 187, 69–79. https://doi.org/10.1677/joe.1.06220

Thistle, R., Cameron, P., Ghorayshi, A., Dennison, L., Scott, K., 2012. Contact Chemoreceptors Mediate Male-Male Repulsion and Male-Female Attraction during Drosophila Courtship. Cell 149, 1140–1151. https://doi.org/10.1016/j.cell.2012.03.045

Toda, H., Zhao, X., Dickson, B.J., 2012. The Drosophila Female Aphrodisiac Pheromone Activates ppk23+ Sensory Neurons to Elicit Male Courtship Behavior. Cell Reports 1, 599–607. https://doi.org/10.1016/j.celrep.2012.05.007

Traynor, K.S., Le Conte, Y., Page, R.E., 2014. Queen and young larval pheromones impact nursing and reproductive physiology of honey bee (Apis mellifera) workers. Behav Ecol Sociobiol 68, 2059–2073. https://doi.org/10.1007/s00265-014-1811-y

Trienens, M., Kraaijeveld, K., Wertheim, B., 2017. Defensive repertoire of *Drosophila* larvae in response to toxic fungi. Mol Ecol 26, 5043–5057. https://doi.org/10.1111/mec.14254

Ulrich, Y., Burns, D., Libbrecht, R., Kronauer, D.J.C., 2016. Ant larvae regulate worker foraging behavior and ovarian activity in a dose-dependent manner. Behav Ecol Sociobiol 70, 1011–1018. https://doi.org/10.1007/s00265-015-2046-2

Umberson, D., Karas Montez, J., 2010. Social Relationships and Health: A Flashpoint for Health Policy. J Health Soc Behav 51, S54–S66. https://doi.org/10.1177/0022146510383501

Uzsák, A., Schal, C., 2013. Sensory Cues Involved in Social Facilitation of Reproduction in Blattella germanica Females. PLoS ONE 8, e55678. https://doi.org/10.1371/journal.pone.0055678

Uzsak, A., Schal, C., 2012. Differential physiological responses of the German cockroach to social interactions during the ovarian cycle. Journal of Experimental Biology 215, 3037–3044. https://doi.org/10.1242/jeb.069997

Vijendravarma, R.K., Narasimha, S., Kawecki, T.J., 2013a. Predatory cannibalism in Drosophila melanogaster larvae. Nat Commun 4, 1789. https://doi.org/10.1038/ncomms2744

Vijendravarma, R.K., Narasimha, S., Kawecki, T.J., 2013b. Predatory cannibalism in Drosophila melanogaster larvae. Nature Communications 4. https://doi.org/10.1038/ncomms2744

Villalta, I., Angulo, E., Devers, S., Cerdá, X., Boulay, R., 2015. Regulation of worker egg laying by larvae in a fission-performing ant. Animal Behaviour 106, 149–156. https://doi.org/10.1016/j.anbehav.2015.05.021

Walker, R.G., 2000. A Drosophila Mechanosensory Transduction Channel. Science 287, 2229–2234. https://doi.org/10.1126/science.287.5461.2229

Wasser, S.K., Barash, D.P., 1983. Reproductive Suppression Among Female Mammals: Implications for Biomedicine and Sexual Selection Theory. The Quarterly Review of Biology 58, 513–538. https://doi.org/10.1086/413545

Wertheim, B., Allemand, R., Vet, L.E.M., Dicke, M., 2006. Effects of aggregation pheromone on individual behaviour and food web interactions: a field study on *Drosophila*: Aggregation pheromone in ecological webs. Ecological Entomology 31, 216–226. https://doi.org/10.1111/j.1365-2311.2006.00757.x

Wertheim, B., Dicke, M., Vet, L.E.M., 2002a. Behavioural plasticity in support of a benefit for aggregation pheromone use in Drosophila melanogaster. Entomologia Experimentalis et Applicata 103, 61–71. https://doi.org/10.1046/j.1570-7458.2002.00954.x

Wertheim, B., Marchais, J., Vet, L.E.M., Dicke, M., 2002b. Allee effect in larval resource exploitation in Drosophila: an interaction among density of adults, larvae, and micro-organisms: Allee effect in larval resource exploitation. Ecological Entomology 27, 608–617. https://doi.org/10.1046/j.1365-2311.2002.00449.x

Wertheim, B., Marchais, J., Vet, L.E.M., Dicke, M., 2002c. Allee effect in larval resource exploitation in Drosophila: an interaction among density of adults, larvae, and micro-organisms. Ecological Entomology 27, 608–617. https://doi.org/10.1046/j.1365-2311.2002.00449.x

White, L.J., Thomson, J.S., Pounder, K.C., Coleman, R.C., Sneddon, L.U., 2017. The impact of social context on behaviour and the recovery from welfare challenges in zebrafish, Danio rerio. Animal Behaviour 132, 189–199. https://doi.org/10.1016/j.anbehav.2017.08.017

White, M.A., Chen, D.S., Wolfner, M.F., 2021. She’s got nerve: roles of octopamine in insect female reproduction. Journal of Neurogenetics 1–22. https://doi.org/10.1080/01677063.2020.1868457

Wicker-Thomas, C., 2007. Pheromonal communication involved in courtship behavior in Diptera. Journal of Insect Physiology 53, 1089–1100. https://doi.org/10.1016/j.jinsphys.2007.07.003

Wijesekera, T.P., Saurabh, S., Dauwalder, B., 2016. Juvenile Hormone Is Required in Adult Males for Drosophila Courtship. PLoS ONE 11, e0151912. https://doi.org/10.1371/journal.pone.0151912

Wilson, E.O., 1971. The insect societies. Cambridge, Mass. : The Belknap Press of Harvard Univ. Press.

Wong, J.W.Y., Meunier, J., Kölliker, M., 2013. The evolution of parental care in insects: the roles of ecology, life history and the social environment: The evolution of parental care in insects. Ecol Entomol 38, 123–137. https://doi.org/10.1111/een.12000

Wu, M., Nern, A., Williamson, W.R., Morimoto, M.M., Reiser, M.B., Card, G.M., Rubin, G.M., 2016. Visual projection neurons in the Drosophila lobula link feature detection to distinct behavioral programs. eLife 5, e21022. https://doi.org/10.7554/eLife.21022

Wyatt, T.D., 2014. Pheromones and animal behavior: chemical signals and signatures, 2nd edition. ed. Cambridge University Press, Cambridge.

Zhou, L., Steller, H., 2003. Distinct Pathways Mediate UV-Induced Apoptosis in Drosophila Embryos. Developmental Cell 4, 599–605. https://doi.org/10.1016/S1534-5807(03)00085-6

Zhu, E.Y., Guntur, A.R., He, R., Stern, U., Yang, C.-H., 2014. Egg-Laying Demand Induces Aversion of UV Light in Drosophila Females. Current Biology 24, 2797–2804. https://doi.org/10.1016/j.cub.2014.09.076

Zhu, Y., 2013. The *Drosophila* visual system: From neural circuits to behavior. Cell Adhesion & Migration 7, 333–344. https://doi.org/10.4161/cam.25521

