## Supplementary figures and table for "Social modulation of oogenesis and egg-laying in *Drosophila melanogaster*"

### Supplementary figure and Tables

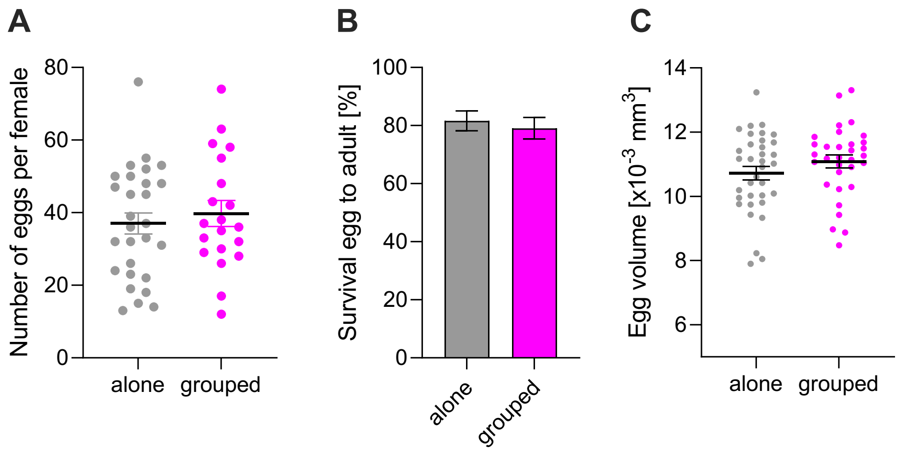


**Supp. Figure S1**: **(A)** Total number of eggs laid in 24h by a wild-type *Oregon-R* mated female alone or in group with 5 mated females laying green fluorescent eggs (GFP). Replicates per group: 20-28. **(B)** Survival of wild-type *Oregon-R* eggs laid by alone or grouped females housed with 5 GFP females into viable adults. Replicates per group: 20-28. **(C)** Average egg volume laid by a wild-type *Oregon-R* mated female alone or in group with 5 other mated females. Replicates per group: 32-35. Error bars indicate standard error of the mean. For full statistical analysis and methods, see **Supplementary Table 1.**

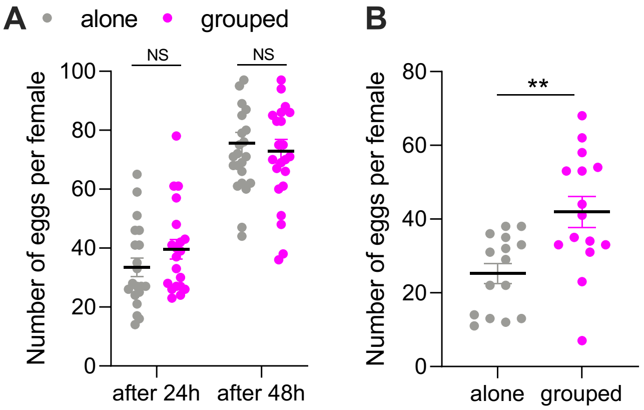


**Supp. Figure S2**: **(A)** Total number of eggs laid in 24h and 48h by a wild-type *Oregon-R* mated female alone or in group with 5 *Oregon-R* males. Replicates per group: 20-24. **(B)** Total number of eggs laid in 24h by a wild-type *Oregon-R* mated female alone or in group with 5 *Oregon-R* males under 24h light conditions. Replicates per group: 15. Error bars indicate standard error of the mean. Stars indicate differences between conditions (*** $p<0.001$, ** $p<0.01$, * $p<0.05$). For full statistical analysis and methods, see **Supplementary Table 1.**

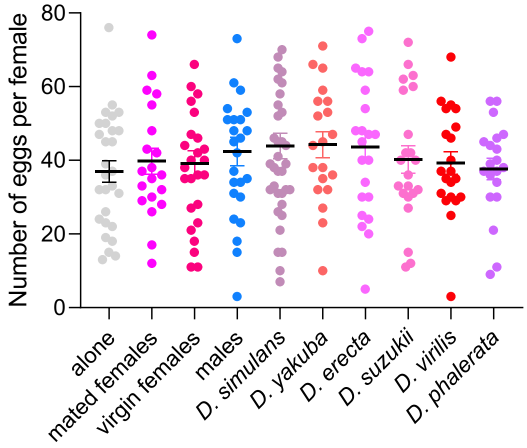


**Supp. Figure S3**: Total number of eggs laid in 24h by a wild-type *Oregon-R* mated female alone or in group with 5 *D. melanogaster* mated females, virgin females, males or with males from different *Drosophila* species. Species were from different relatedness and were ranged from the least distant to the most distant related one from *D. melanogaster*: *D. simulans, D. yakuba, D.erecta, D. suzukii, D. virilise, D. phalerata*. Replicates per group: 20-37. Error bars indicate standard error of the mean. For full statistical analysis and methods, see **Supplementary Table 1.**

**Supp. Table S1**: Summary of performed statistical analyses.

| Fig | Test | Response variable | Explanatory factor | Results | Post-hoc test (only p-value <0.05) |
|---|---|---|---|---|---|
| 1A | Kruskal-Wallis | Egg-laying start-time | Group condition [alone, grouped with 5 GFP, grouped with 5 RFP] | $\chi^2 = 28.172$, df = 2, $p < 0.001$ | Alone vs 5GFP: $p < 0.001$, Alone vs 5RFP: $p < 0.001$ |
| 1B | Kruskal- Wallis | Egg-laying start-time | Group size [1, 2, 3, 4, 5, 6, 12, 18, 24, 50] | $\chi^2 = 155.59$, df = 10, $p < 0.001$ | NA |
| 1C | Kruskal- Wallis | Survival egg to adult at 100% food | Group size [1, 6, 12, 24, 50] | $\chi^2 = 12.679$, df = 4, $p = 0.01$ | 1 vs 12: $p = 0.004$, 1 vs 24: $p = 0.019$ |
| | | Survival egg to adult at 50% food | | $\chi^2 = 35.198$, df = 4, $p < 0.001$ | 1 vs 24: $p = 0.013$, 1 vs 50: $p < 0.001$, 12 vs 50: $p < 0.001$, 24 vs 50: $p = 0.005$, 50 vs 6: $p < 0.001$ |
| | | Survival egg to adult at 25% food | | $\chi^2 = 68.475$, df = 4, $p < 0.001$ | 1 vs 12: $p < 0.001$, 1 vs 24: $p < 0.001$, 1 vs 50: $p < 0.001$, 1 vs 6: $p < 0.001$, 12 vs 24: $p = 0.033$, 12 vs 50: $p < 0.001$, 24 vs 50: $p < 0.001$, 24 vs 6: $p < 0.001$, 50 vs 6: $p < 0.001$ |
| | | Survival egg to adult at 10% food | | $\chi^2 = 39.489$, df = 4, $p < 0.001$ | 1 vs 12: $p < 0.001$, 1 vs 24: $p < 0.001$, 1 vs 50: $p < 0.001$, 1 vs 6: $p = 0.009$, 24 vs 6: $p < 0.001$, 50 vs 6: $p = 0.001$ |
| 1D | Binomial Zero-inflated linear model | Survival egg to adult | Group condition [host and guest homog groups, group guest in synch with host (g1), group guest out of synch with host (g2), alone guest out of synch with host (a)] | $\chi^2 = 131.57$, df = -14, $p < 0.001$ | Homog guest vs a guest: $p < 0.001$, homog guest vs g2 guest: $p = 0.0014$, a guest vs g1 guest: $p < 0.001$, g1 guest vs g2 guest: $p < 0.001$, homog host vs a host: $p = 0.01$ |
| $1E_1$ | Kruskal- Wallis | Latency to leave the initial dish | Social condition [grouped-alone, alone-alone, alone-grouped] | $\chi^2 = 16.990$, df = 2, $p < 0.001$ | Grouped-alone vs alone-alone: $p < 0.001$, grouped-alone vs alone-grouped: $p < 0.001$ |
| $1E_2$ | Poisson Generalized linear model | Number of tunnel crosses | Social condition [grouped-alone, alone-alone, alone-grouped] | $F = 5.107$, $p = 0.009$ | Grouped-alone vs alone-alone: $p = 0.007$ |
| $1E_3$ | Binomial Generalized linear model | Index egg preference | Social condition [grouped-alone, alone-alone, alone-grouped] | $F = 15.97$, $p < 0.001$ | Grouped-alone vs alone-alone: $p = 0.025$, grouped-alone vs alone-grouped: $p = 0.001$ |
| | One-sample Wilcoxon | Index egg preference | Comparison for each social condition to 0 (no preference) | NA | Grouped-alone vs 0: $V = 209$, $p < 0.001$,<br>alone-grouped vs 0: $V = 81$, $p = 0.05$ |
| 2A | Two-way repeated- | Cumulative number of eggs | Circadian time x Group [alone, grouped] | $F(24,1080) = 12.83$, df = 24, $p < 0.001$ | Alone vs grouped CT11: $p < 0.001$, Alone vs grouped CT12: $p < 0.001$, Alone vs grouped CT12 day 2: $p = 0.029$ |

| | | | | | |
|---|---|---|---|---|---|
| | measurements Anova | | | | |
| 2B | Two-way repeated-measurements Anova | Cumulative number of eggs | Circadian time x Group [alone, grouped] | F(24,912) = 7.532, df = 24, p < 0.001 | Alone vs grouped CT9: p = 0.041, Alone vs grouped CT10: p < 0.001, Alone vs grouped CT11: p < 0.001, Alone vs grouped CT12: p < 0.001, Alone vs grouped CT13: p = 0.009 |
| 2C | Two-way repeated-measurements Anova | Cumulative number of eggs | Circadian time x Group [alone, grouped] | F(24,672) = 1.213, df = 24, p = 0.221 | No p-value < 0.05 |
| 2D | Two-way repeated-measurements Anova | Cumulative number of eggs | Circadian time x Group [alone, grouped] | F(24,672) = 13.97, df = 24, p < 0.001 | Alone vs grouped CT8: p = 0.044, Alone vs grouped CT9: p = 0.044, Alone vs grouped CT10: p = 0.002, Alone vs grouped CT11: p < 0.001, Alone vs grouped CT12: p < 0.001, Alone vs grouped CT13: p < 0.001, Alone vs grouped CT14: p < 0.001, Alone vs grouped CT15: p < 0.001, Alone vs grouped CT16: p < 0.001, Alone vs grouped CT17: p = 0.001, Alone vs grouped CT18: p = 0.002, Alone vs grouped CT19: p = 0.004, Alone vs grouped CT20: p = 0.005, Alone vs grouped CT21: p = 0.008, Alone vs grouped CT22: p = 0.014, Alone vs grouped CT23: p = 0.020, Alone vs grouped CT0: p = 0.033, Alone vs grouped CT1: p = 0.044, Alone vs grouped CT2: p = 0.050, Alone vs grouped CT3: p = 0.050, Alone vs grouped CT4: p = 0.048 |
| 2E | Type 3 two-way Anova | Egg-laying start-time | Group [alone, group] x Light [LL, DD, LD] | F = 57.981, p < 0.001 | DD-LD alone: p < 0.001, DD-LL alone: p < 0.001, LD-LL alone: p < 0.001, alone-group LD: p < 0.001, alone-group LL: p < 0.001 |
| 2F | Two-way repeated-measurements Anova | Cumulative number of eggs | Alone CT [CT2, CT5] | F(24,912) = 4.571, df = 24, p < 0.001 | CT10 vs CT13: p < 0.001, CT11 vs CT14: p < 0.001, CT2 vs CT15: p = 0.035 |
| | | | Grouped CT [CT2, CT5] | F(24,888) = 2.142, df = 24, p = 0.001 | No p-value < 0.05 |
| 2G | Type 3 two-way Anova | Egg-laying start-time | Group [alone, grouped] x Circadian time | F = 4.185, p = 0.044 | Alone CT2 vs alone CT5: p < 0.001, group CT2 vs group CT5: p = 0.042, CT2 alone vs CT2 group: p < 0.001, CT5 alone vs CT5 group: p < 0.001 |
| 3A | Kruskal-Wallis | Egg-laying start-time | Group condition [alone, mated, virgin, males, fru-] | $\chi^2$ = 32.815, df = 4, p < 0.001 | Alone vs fru-: p = 0.002, alone vs males: p < 0.001, alone vs mated: p < 0.001, alone vs virgins: p < 0.001 |

| | | | | | |
|---|---|---|---|---|---|
| 3B | Kruskal-Wallis | Egg-laying start-time | Group condition [alone, mel, sim, yak, er, suz, vir, pha] | $\chi^2$ = 67.915, df = 7, p < 0.001 | Alone vs er: p < 0.001, alone vs mel: p < 0.001, alone vs pha: p = 0.002, alone vs sim: p < 0.001, alone vs suz: p < 0.001, alone vs vir: p < 0.001, alone vs yak: p < 0.001 |
| 3C | Kruskal-Wallis | Egg-laying start-time | Group condition [alone, ctrl females, ctrl males, oe females, oe males] | $\chi^2$ = 30.452, df = 4, p < 0.001 | Ctrl females vs alone: p < 0.001, ctrl males vs alone: p < 0.001, oe females vs alone: p < 0.001, oe males vs alone: p = 0.003 |
| 3D | Wilcoxon-Mann-Whitney | Egg-laying start-time of WT | Group condition [alone, grouped] | W = 564, p < 0.001 | NA |
| | | Egg-laying start-time of $ort^{1}$ | | W = 193, p = 0.944 | NA |
| | | Egg-laying start-time of $piezo^{KO}$ | | W = 628, p < 0.001 | NA |
| | | Egg-laying start-time of nompC | | W = 414, p < 0.001 | NA |
| | | Egg-laying start-time of $ppk23^{-}$ | | W = 501, p < 0.001 | NA |
| | | Egg-laying start-time of $ppk29^{-}$ | | W = 448, p < 0.001 | NA |
| | | Egg-laying start-time of $orco^{1}$ | | W = 515, p < 0.001 | NA |
| 3E | Kruskal-Wallis | Egg-laying start-time | Group condition [alone, 1 female + mirror, 2 females] | $\chi^2$ = 14.739, df = 2, p < 0.001 | 1 fem vs 1 fem + mirror: p = 0.016, 1 fem vs 2 fem: p < 0.001 |
| 4A | Type 3 two-way Anova | Egg-laying start-time | Group [alone, grouped] x Petri dish size [small (0.64 fly/$cm^2$), medium (0.24 fly/$cm^2$), big (0.15 fly/$cm^2$), huge (0.066 fly/$cm^2$)] | F = 6.629, p < 0.001 | Big vs small grouped: p < 0.001, huge vs medium grouped: p = 0.002, huge vs small grouped: p < 0.001, medium vs small grouped: p = 0.004, alone vs grouped big: p = 0.038, alone vs grouped huge: p = 0.038, alone vs grouped medium: p < 0.001, alone vs grouped: p < 0.001 |
| 4B | Linear correlation | Egg-laying start-time<br>Density | NA | F = 7.289, $R^2$ = 0.785, p = 0.114 | NA |
| 4C | Linear correlation | Number of contacts<br>Density | NA | F = 4714, $R^2$ = 0.999, p = 0.009 | NA |
| 4D | Linear correlation | Inter-individual distance<br>Density | NA | F = 4.954, $R^2$ = 0.832, p = 0.269 | NA |
| 4E | Kruskal-Wallis | Egg-laying start-time | Group condition [alone t4t5xkir, group t4t5xkir, alone t4t5xor, group | $\chi^2$ = 65.265, df = 5, p < 0.001 | Alone t4t5xkir vs group t4t5xor : p < 0.001, alone t4t5xkir vs group kirxor : p < 0.001, alone t4t5xor vs group t4t5xor : p < 0.001, alone t4t5xor vs group kirxor: p < 0.001, group t4t5xor: |

| | | | | | |
|---|---|---|---|---|---|
| | | | t4t5xor, alone kirxor, group kirxor] | | p < 0.001, group t4t5xor vs group t4t5xkir: p < 0.001, alone kirxor vs group kirxor: p < 0.001, group kirxor vs group t4t5xkir: p< 0.001 |
| 4F | Kruskal-Wallis | Egg-laying start-time | Group condition [alone lc11xkir, group lc11xkir , alone lc11xor, group lc11xor, alone kirxor, group kirxor] | $\chi^2 = 73.514$, df = 5, p < 0.001 | Alone lc11xkir vs group lc11xor: p < 0.001, alone lc11xkir vs group kirxor: p < 0.001, alone lc11xkir vs group lc11xkir: p < 0.001, alone lc11xor vs group lc11xor: p < 0.001, alone lc11xor vs group kirxor: p < 0.001, alone lc11xor vs group lc11xkir: p = 0.002, group lc11xor vs alone kirxor: p < 0.001, alone kirxor vs group kirxor: p < 0.001, alone kirxor vs group lc11xkir: p < 0.001 |
| 4G | Kruskal-Wallis | Egg-laying start-time | Group condition [alone lc10xkir, group lc10xkir, alone lc10xor, group lc10xor, alone kirxor, group kirxor] | $\chi^2 = 68.334$, df = 5, p < 0.001 | Alone lc10xkir vs group lc10xkir: p < 0.001, alone lc10xkir vs group lc10xor: p < 0.001, alone lc10xkir vs group kirxor: p <0.001, alone lc10xor vs group lc10xkir: p < 0.001, alone lc10xor vs group lc10xor: p < 0.001, alone lc10xor vs group kirxor: p < 0.001, alone kirxor vs group lc10xkir: p < 0.001, alone kirxor vs group lc10xor: p < 0.001, alone kirxor vs group kirxor: p < 0.001, group lc10xkir vs group kirxor: p = 0.023 |
| 5A | Wilcoxon-Mann-Whitney | Ovulation rate 2h under light | Group condition [alone, grouped] | W = 245, p = 0.013 | NA |
| | | Ovulation rate 6h under light | | W = 624.5, p = 0.025 | NA |
| | | Ovulation rate 2h under dark | | W = 540.5, p = 0.829 | NA |
| 5B | Wilcoxon-Mann-Whitney | Number of s14 oocytes 2h under light | Group condition [alone, grouped] | W = 906, p = 0.004 | NA |
| | | Number of s14 oocytes 6h under light | | W = 2333, p < 0.001 | NA |
| | | Number of s14 oocytes 2h under dark | | W = 1334, p = 0.801 | NA |
| 5E | Wilcoxon-Mann-Whitney | Number of mature eggs 2h under light | Group condition [alone, grouped] | W = 219.5, p = 0.012 | NA |
| | | Number of mature eggs 6h under light | | W = 400, p = 0.544 | NA |
| | | Number of mature eggs 2h under dark | | W = 249.5, p = 0.758 | NA |

| | | | | | |
|---|---|---|---|---|---|
| 6A | Kruskal-Wallis | Number of s14 oocytes | Group condition [alone, alone + methoprene, grouped] | $\chi^2 = 214.389$, df = 2, p < 0.001 | Alone vs grouped: p = 0.002, alone vs alone + meth: p < 0.001, grouped vs alone + meth: p = 0.954 |
| 6B | Kruskal-Wallis | Egg-laying start-time | Group condition [alone, alone + methoprene, grouped] | $\chi^2 = 33.480$, df = 2, p < 0.001 | Alone vs grouped: p < 0.001, alone vs alone + meth: p = 0.816, grouped vs alone + meth: p < 0.001 |
| 6C | Kruskal-Wallis | Ovulation rate | Group condition [alone, alone + methoprene, grouped] | $\chi^2 = 10.167$, df = 2, p = 0.006 | Alone vs grouped: p = 0.01, alone vs alone + meth: p = 1, grouped vs alone + meth: p = 0.007 |
| 6D | Anova | Number of s14 oocytes | Grouped condition of ort[1] [alone, grouped + methoprene, grouped] | F = 9.899, df = 2, p < 0.001 | Alone vs grouped: p = 0.974, alone vs meth: p < 0.001, grouped vs meth: p < 0.001 |
| | Wilcoxon-Mann-Whitney | Number of s14 oocytes | Grouped condition of OR [alone, grouped] | W = 1328.5, p < 0.001 | NA |
| 6E | Kruskal-Wallis | Egg-laying start-time | Group condition [alone, grouped + methoprene, grouped] | $\chi^2 = 48$, df = 29, p = 0.015 | Alone vs grouped: p = 1, alone vs meth: p = 0.974, grouped vs meth: p = 0.932 |
| S1A | Student's t-test | Number of eggs per female | Group condition [alone, grouped] | t = -0.600, df = 40.675, p = 0.551 | NA |
| S1B | Wilcoxon-Mann-Whitney | Survival egg to adult | Group condition [alone, grouped] | W = 279.5, p = 0.540 | NA |
| S1C | Wilcoxon-Mann-Whitney | Egg volume | Group condition [alone, grouped] | W = 477, p = 0.302 | NA |
| S2A | Wilcoxon-Mann-Whitney | Number of eggs after 24h | Group condition [alone, grouped] | W = 152.5, p = 0.203 | NA |
| | Student's t-test | Number of eggs after 48h | Group condition [alone, grouped] | t = 0.495, df = 44.461, p = 0.623 | NA |
| S2B | Student's t-test | Number of eggs per female | Group condition [alone, grouped] | t = -3.343, df = 23.809, p = 0.003 | NA |
| S3 | Anova | Number of eggs per female | Group condition [alone, mated, virgins, males, sim, yak, er, suz, vir, pha] | F = 0.699, df = 9, p = 0.571 | No p-value < 0.05 |
